# A giant cell enhancer achieves cell-type specificity through activation via TCP and repression by Dof transcription factors

**DOI:** 10.1101/2020.08.28.270454

**Authors:** Lilan Hong, Clint S. Ko, S. Earl Kang, Jose L. Pruneda-Paz, Adrienne H. K. Roeder

## Abstract

Proper pattern formation relies on the tight coordination of cell fate specification and cell cycle regulation in growing tissues. How this can be organized at enhancers that activate gene expression necessary for differentiation is not well understood. One such example is the patterning of the *Arabidopsis thaliana* sepal epidermis where giant cell fate specification is associated with the endoreduplication cell cycle. Previously, we identified an enhancer region capable of driving giant cell-specific expression. In this study, we use the giant cell enhancer as a model to understand the regulatory logic that promotes cell-type specific expression. Our dissection of the enhancer revealed that giant cell specificity is achieved primarily through the combination of two elements: an activator and a repressor. TCP transcription factors are involved in activation of non-specific expression throughout the epidermis with higher expression in endoreduplicated giant cells than small cells. Dof transcription factors act via the second element to repress activity of the enhancer and limit expression to giant cells. Thus, we find that cell-type specific expression emerges from the combined activities of two broadly acting enhancer elements.

## Introduction

Plant and animal organs are composed of different types of cells that perform specialized roles. Cell-type specification is driven by the establishment of distinct patterns of gene expression, as has been confirmed recently by single-cell RNA-seq (Brady et al., 2007; Denyer et al., 2019; Jaitin et al., 2014; Lee and Schiefelbein, 2002; Ryu et al., 2019). Cell-type specific expression patterns are generally orchestrated through transcription factors binding to enhancers, short non-coding DNA sequences (Long et al., 2016). Enhancers typically contain clusters of transcription factor binding sites, acting as platforms to integrate information (Buecker and Wysocka, 2012). The spatial and temporal specificity of transcriptional regulation through enhancers has been extensively studied in several model organisms, including mouse, sea urchins, chicks, and fruit flies (Davidson, 2010; Spitz and Furlong, 2012). Recently, many enhancers have been identified in *Arabidopsis thaliana* (henceforth Arabidopsis) and other plants through whole genome methods, including hypersensitivity to DNase I (Oka et al., 2017; Yan et al., 2019; Zhu et al., 2015) and ATAC-seq (Ricci et al., 2019; Tannenbaum et al., 2018). However, relatively little is known about how individual enhancers function mechanistically in plants to generate cell-type specific expression patterns.

To elucidate the regulatory logic through which an individual enhancer drives cell-type specific expression, we used the Arabidopsis sepal giant cell model system. Sepals, the outermost floral organ, protect the inner androecium and gynoecium during floral development. On the abaxial epidermis of Arabidopsis sepals, epidermal pavement cells form patterns of two cell types: mitotically dividing small cells and elongated, endoreduplicating giant cells, which scatter between the small cells (Figure 1B) (Roeder et al., 2010). Endoreduplication is a specialized cell cycle wherein cells continue to grow and replicate their DNA but fail to undergo mitosis, generating enlarged, polyploid cells (Traas et al., 1998). Thus, giant cells are a good model system for understanding the intersection of cell-type specification with cell division and growth. Giant cell fate specification and differentiation is promoted by the epidermal specification pathway, including Homeodomain-leucine zipper (HD-ZIP) Class IV transcription factors Arabidopsis thaliana MERISTEM LAYER 1 (ATML1) and HOMEODOMAIN GLABROUS 11 (HDG11), as well as intercellular signaling proteins DEFECTIVE KERNEL 1 (DEK1) and Arabidopsis CRINKLY 4 (ACR4) (Roeder et al., 2012). The endomembrane trafficking protein SEC24A restricts the formation of giant cells by modulating this pathway (Qu et al., 2014). Downstream of the epidermal specification pathway, the SIAMESE family Cyclin Dependent Kinase Inhibitor (CKI) LOSS OF GIANT CELLS FROM ORGANS (LGO, also known as SMR1) is required for endoreduplication of giant cells (Meyer et al., 2017; Roeder et al., 2010; 2012; Schwarz and Roeder, 2016). Giant cells affect sepal curvature (Roeder et al., 2012) and are hypothesized to play a role in defense against pathogens based on transcriptomic analysis (Schwarz and Roeder, 2016). Stochasticity in gene expression appears to be important for initiating the scattered pattern of giant cells between smaller cells. Fluctuation of ATML1 expression to a high level during the G2 phase of the cell cycle specifies giant cell fate (Meyer et al., 2017).

**Figure 1.**
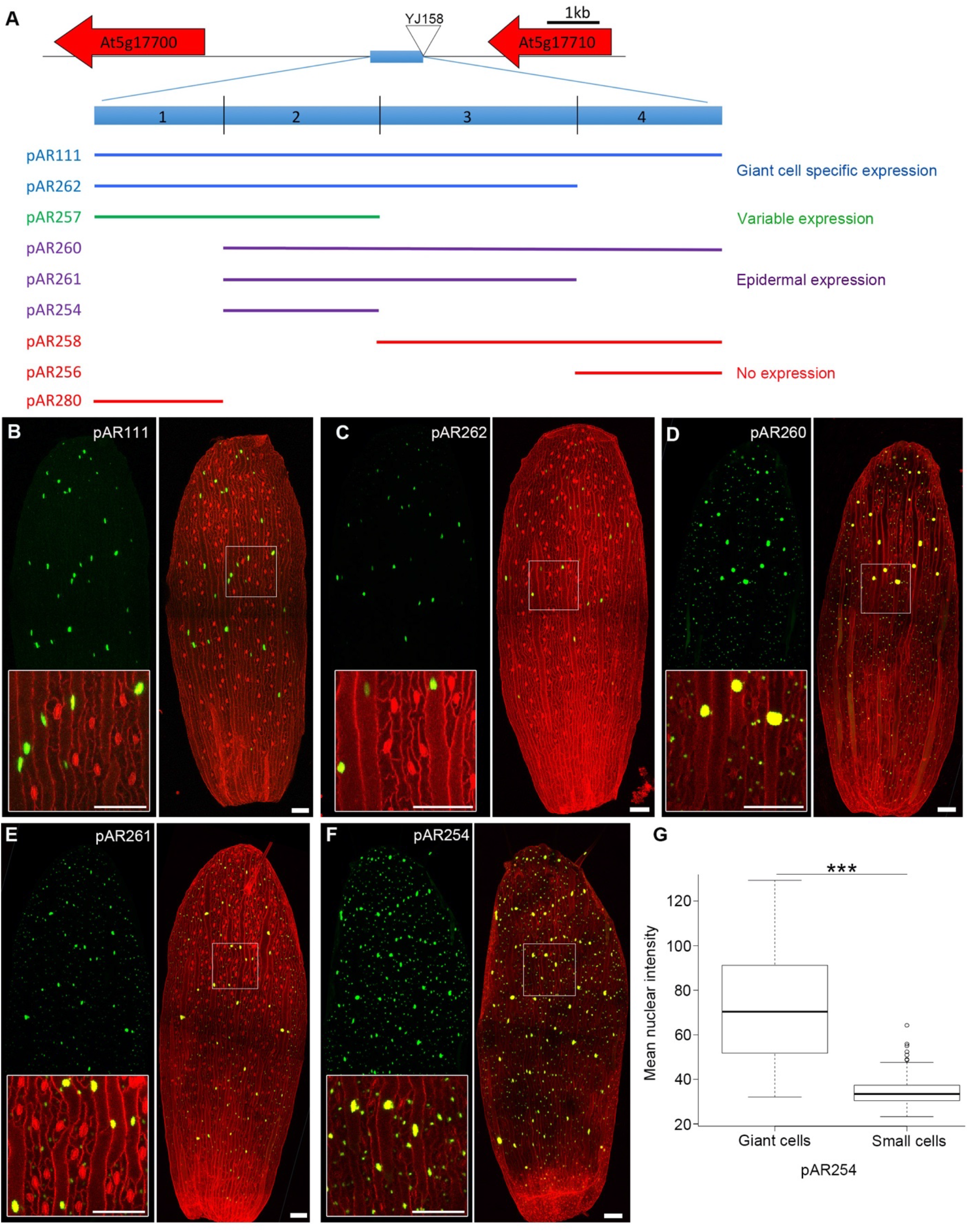
Dissection of 1kb giant cell enhancer. **(A)** The giant cell enhancer is a 1 kb sequence 3.2 kb upstream of *AT5G17700*, a gene encoding a MATE efflux family protein. The 1 kb enhancer was originally identified because it flanked the YJ158 enhancer trap T-DNA insertion (triangle), which produced the same giant cell-specific expression pattern. The enhancer was divided into four regions. Different constructs containing different regions of the enhancer for testing enhancer elements are diagrammed below. Reporter lines with these constructs were tested for their ability to produce expression of the nuclear localized 3×Venus-N7 reporter under a -60 minimal 35S promoter. Enhancer fragments in the different reporter lines are color coded with their expression patterns: blue denotes giant cell-specific expression, purple for epidermal expression, green for expression patterns that vary between giant cell-specific and broad epidermal independent T1 lines (further classified in Figure 2), and red for no expression. **(B-F)** Confocal images of stage 14 sepals from different reporter lines. Images on the left, 3×Venus (green) marking the nuclei of cells expressing the reporter; Images on the right, 3×Venus (green) signals merged with propidium iodide (PI, red) stained cell wall signals. Insets overlapping the base of the green channel images, show magnified views of the cells outlined with the box. Expression of the 3×Venus is restricted to giant cells in reporter lines *pAR111* and *pAR262* (B,C). The reporter has a broader expression in *pAR260, pAR261* and *pAR254* (D,E,F). Note that stomatal guard cells stain as red oblongs. Scale bars: 100 μm. **(G)** Mean fluorescent intensity of the 3×Venus reporter in giant cell nuclei versus small cell nuclei of pAR254 sepals, which is expressed in a broad epidermal pattern. Note that the expression (intensity) is about 2-fold higher in giant cell nuclei. The boxes extend from the lower to upper quartile values of the data, with a line at the median. The whiskers extend past 1.5 of the interquartile range. ***p < 0.001, significant difference by two tailed t-test, p = 4.29E-13; n = 210 small cells and 48 giant cells. See also Figures S1 and S2.

Our previous work described an enhancer region capable of driving giant cell-specific expression in sepals (Roeder et al., 2012). The region was identified based on an enhancer trap line (YJ158) in which the T-DNA was inserted 4.7 kb upstream of *AT5G17700* (Figure 1A) (Eshed et al., 2004). *AT5G17700* encodes a MATE efflux family protein. The giant cell-specific enhancer (hereafter referred to as the giant cell enhancer) was functionally defined as the 1024 bp region immediately upstream of the enhancer trap T-DNA insertion; this region was sufficient for giant-cell-specific expression in either orientation (Figure 1A) (Roeder et al., 2012). Expression of a reporter driven by the enhancer is regulated by ATML1 and the other members of the giant cell specification pathway (Roeder et al., 2012). In the leaf, the giant cell enhancer activates reporter expression both in giant epidermal pavement cells and leaf margin cells (Eshed et al., 2004; Roeder et al., 2012). The full 4.7 kb upstream region of *AT5G17700* that includes the giant cell enhancer drives reporter expression in giant cells of young sepals as well as in other cell types (e.g. cells in petals and the style), suggesting that the giant cell enhancer is part of a larger regulatory region for *AT5G17700* (Roeder et al., 2012).

Here we dissected the 1 kb giant cell enhancer into smaller regions and revealed that the enhancer is comprised of an element promoting broad epidermal expression and a region that limits this expression to highly endoreduplicated giant cells in the sepal. A genome-wide yeast one-hybrid screen for transcription factors binding to the enhancer fragments identified Teosinte branched1, Cycloidea, Proliferating cell factor (TCP) and DNA-binding one zinc finger (Dof) transcription factors, among others. We show that TCP transcription factors are involved in activating a broad epidermal pattern of expression. The Dof transcription factors broadly repress expression, leading to giant cell-specific expression.

## Results

### Dissection of the giant cell enhancer delineates regions that drive broad epidermal expression and limit expression to giant cells

To identify regulatory modules, the 1024 bp giant cell enhancer was dissected into four regions: 1 to 208 bp (Region 1), 209 to 449 bp (Region 2), 450 to 760 bp (Region 3), and 761 to 1024 bp (Region 4) (Figure 1A, and S1). Fragments containing these regions or combinations thereof were cloned into a plasmid containing a minimal 35S promoter (mini35S) driving a nuclear-localized yellow fluorescent reporter (3×Venus-N7) (Roeder, et al. 2012). Patterns of Venus reporter expression were characterized in T1 transgenic lines from different constructs to assess the capabilities of the different enhancer fragments to activate transcription. The full 1024 bp enhancer (Region 1-2-3-4, contained in *pAR111*) activated expression specifically in giant cells in the sepal (Figure 1B), as previously shown (Roeder et al., 2012). Region 1 alone (in *pAR280*), Region 4 alone (in *pAR256*), or Regions 3 and 4 together (Region 3-4, in *pAR258*) did not activate reporter expression (Figure 1A and Figure S2). Region 2 alone (in *pAR254*), Region 2 with Region 3 (Region 2-3, in *pAR261*), and Region 2 with Regions 3 and 4 (Region 2-3-4, in *pAR260*) all drove ubiquitous expression in the epidermis, not specific to giant cells (Figure 1A, D-F). Although the reporter was expressed in all epidermal pavement cells for this broad epidermal expression pattern, the expression level was about 2 fold higher in giant cells than small cells, as quantified by mean fluorescence intensity of the reporter in the nucleus (Figure 1G). Regions 1, 2, and 3 together (Region 1-2-3, in *pAR262*) were sufficient for driving reporter expression specifically in endoreduplicated giant cells (Figure 1C).

Regions 1 and 2 together (Region 1-2, in *pAR257*) drove a varied expression pattern in independent T1 transgenic lines, among which the majority showed giant cell specificity. To further characterize the expression specificity of Region 1-2, we classified the expression pattern of these T1 transgenic plants into three categories: (1) giant cell specific (type I) (Figure 2A); (2) enhanced signal in giant cells with additional signal in the small cells at the sepal tip (type II, giant cell enhanced) (Figure 2B); (3) broad epidermal signal (type III) (Figure 2C). We quantified the percentage of *pAR257* (Region 1-2) independent T1 transgenic plants showing these three expression patterns (47.8% giant cell specific, 34.8% giant cell enhanced, and 17.4% epidermal), and compared it with those of *pAR111* (Region 1-2-3-4), *pAR254* (Region 2) and *pAR261* (Region 2-3) plants. *pAR111* (Region 1-2-3-4) transgenic plants all showed giant cell specific signal. For *pAR254* (Region 2) and *pAR261* (Region 2-3), the majority of the lines had type III, broad epidermal signal, although occasionally, they showed type II, giant cell enhanced expression pattern (Figure 2D). Together, these data suggest that Region 1-2 play an important role in generating the giant cell specificity of enhancer activity. Region 2 is sufficient to produce broad epidermal expression, while Region 1 is required for limiting expression to giant cells. In addition, Region 3 and 4 play a small role in enhancing giant cell specificity when present in combination with Region 1; therefore, we focused our further analysis on Regions 1 and 2.

**Figure 2.**
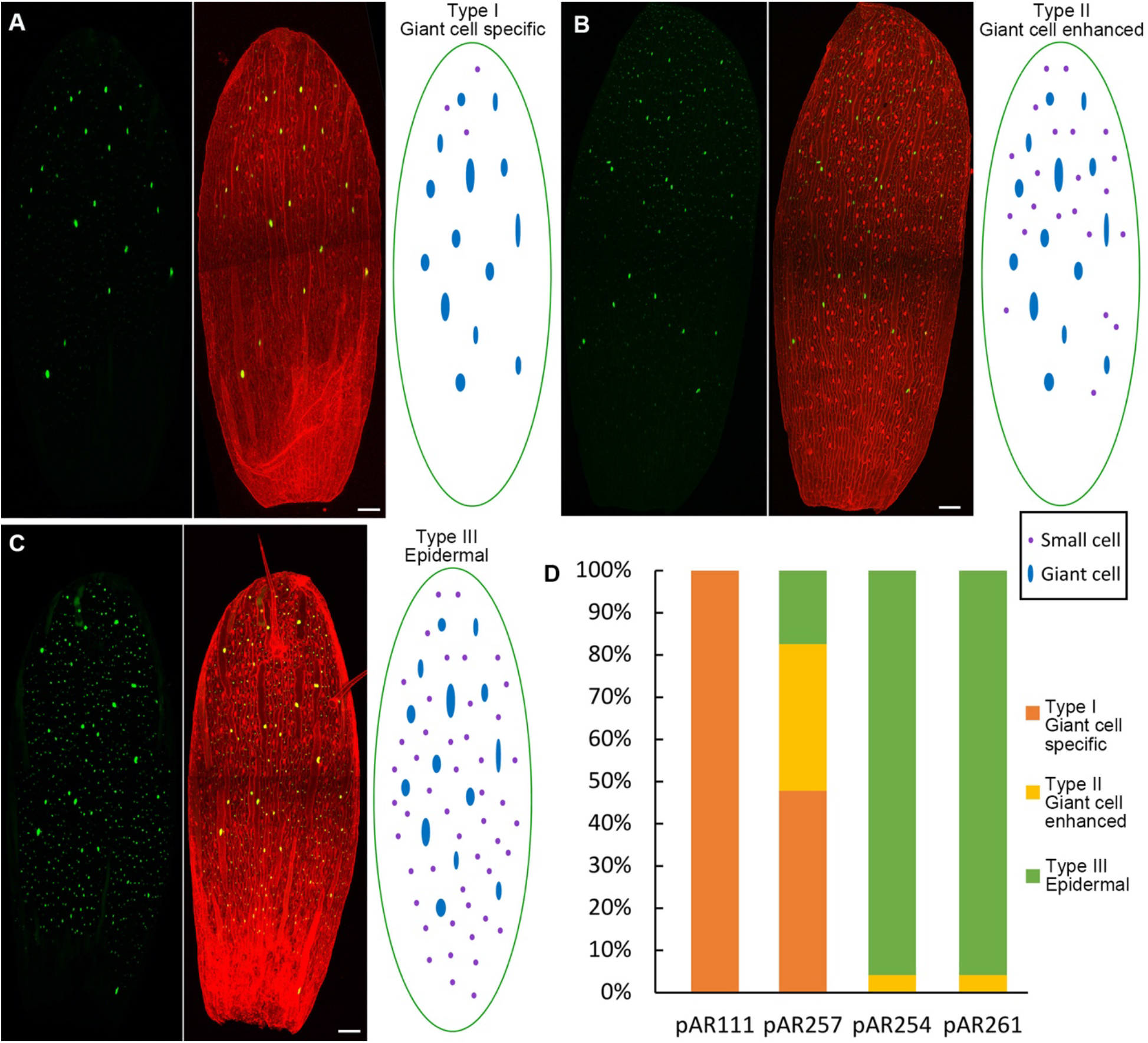
The reporter lines show three types of expression pattern. **(A-C)** The three types of expression pattern detected from reporter lines with *pAR257* (Region 1-2). Images on the left, 3×Venus (green) marking the nuclei of cells expressing the reporter; images in the middle, 3×Venus (green) signals merged with PI (red) stained cell wall signals; images on the right, schematic summarizing the pattern. In type I (giant cell specific) expression pattern, 3×Venus is expressed mainly in giant cells (A). In type II (giant cell enhanced) expression pattern, 3×Venus is expressed both in giant cells and in small cells located in the tip part (B). In type III (epidermal) expression pattern, 3×Venus is expressed broadly throughout the epidermis (C). Scale bars: 100 μm. **(D)** Percentages of the three types of reporter expression patterns observed across different reporter lines. *pAR111* (Region 1-2-3-4) sepals all show the type I pattern. *pAR254* (Region 2) and *pAR261* (Region 2-3) have predominantly type III expression pattern.

### TCP transcription factors promote activity of the enhancer

To identify transcription factors interacting with Regions 1 and 2 of the enhancer, we performed a full-genome yeast one-hybrid screen with a library of 1956 transcription factors. We analyzed Region 1, Region 2, and a 153 bp region overlapping the junction of these regions separately (Figure S1); the overlap was intended to catch interactions that might be disrupted at the boundary between Region 1 and Region 2. We identified 111 high-confidence (Table 1) and 43 low-confidence interactions (Table S1) in total for all three assays (Datasets 1-3). We paid particular attention to transcription factors that specifically bound one region and not the other because they might play roles in driving the distinct expression patterns conferred by these regions.

**Table 1.**
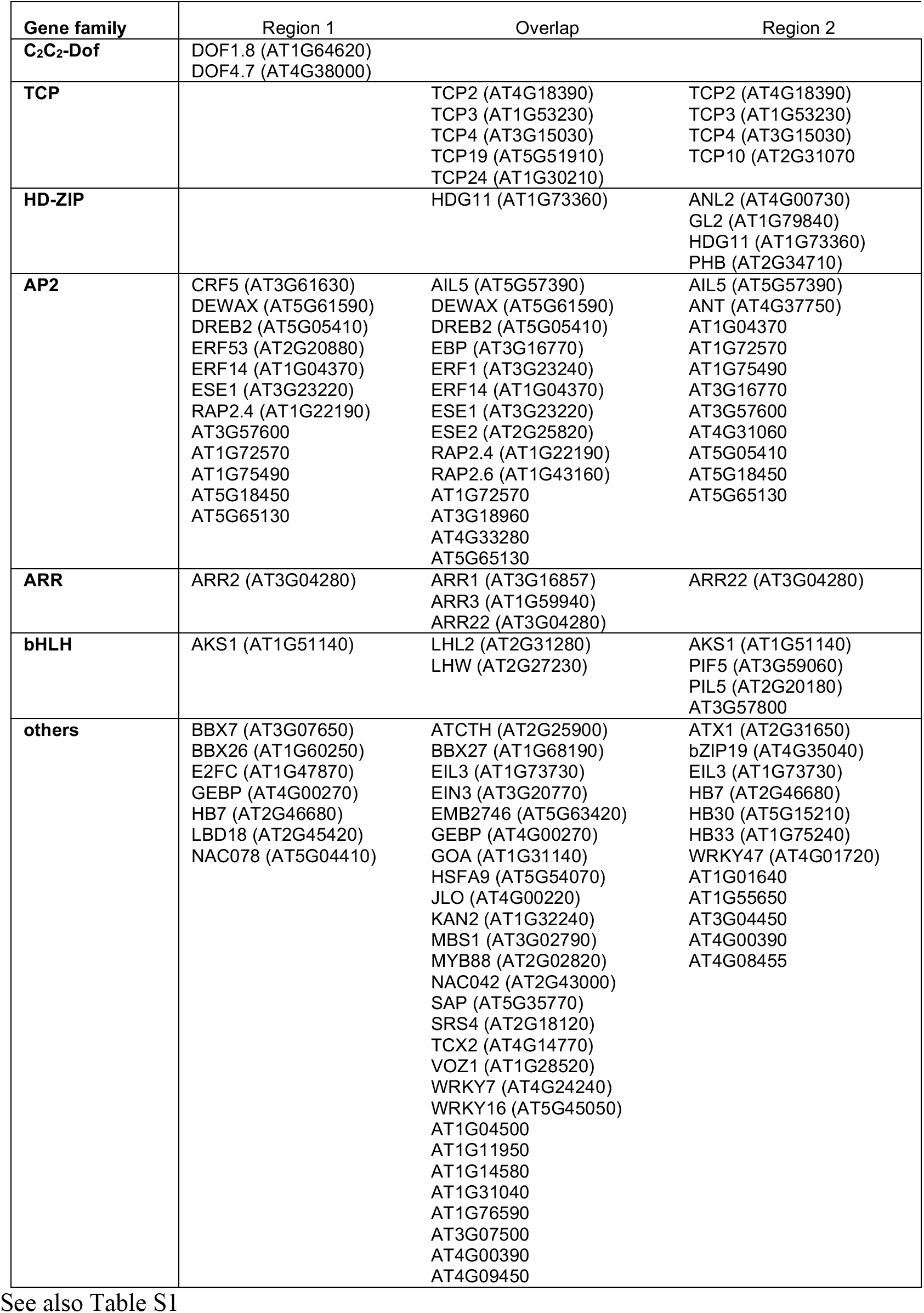
Full-genome yeast one-hybrid screen results: high-confidence interactions.

In the screen, several Class II CINCINNATA-like TCP transcription factors (TCP2, TCP3, TCP4, TCP10) interacted with Region 2 and the overlap of the enhancer, but not Region 1. To test for regulation of the enhancer by Class II TCPs, we crossed the full 1kb enhancer reporter (*pAR111*) to the *jaw-1D* mutant (Palatnik et al., 2003). The *jaw-1D* line overexpresses miR319a, a microRNA that targets *TCP2, TCP3, TCP4, TCP10*, and *TCP24*, thus leading to downregulation of these *TCP*s (Palatnik et al., 2003). In *pAR111* and *jaw-1D* F1 heterozygotes, sepals had fewer cells expressing the reporter (Figure 3B and 3C), compared with sepals from *pAR111* and wild-type (Col-0) F1 heterozygotes (Figure 3A and 3C). Knockdown of TCPs only affected the expression of the pAR111 reporter and not the presence of giant cells in the sepal, as identified by morphology (Figure 3B, PI stain). In *pAR111 × jaw-1D* F1 hybrid sepals, the number of giant cells was similar to that in *pAR111* × wild-type F1 sepals (18.6 ± 2.3, mean ± SD, n = 5 in *pAR111 × jaw-1D* F1; 17 ± 4.1, mean ± SD, n = 5 in *pAR111* × wild-type F1; non-significant difference by t test); however, in *jaw-1D × pAR111* F1 sepals, most of these giant cells were not detectably expressing the *pAR111* reporter (Figure 3B and 3C).

**Figure 3.**
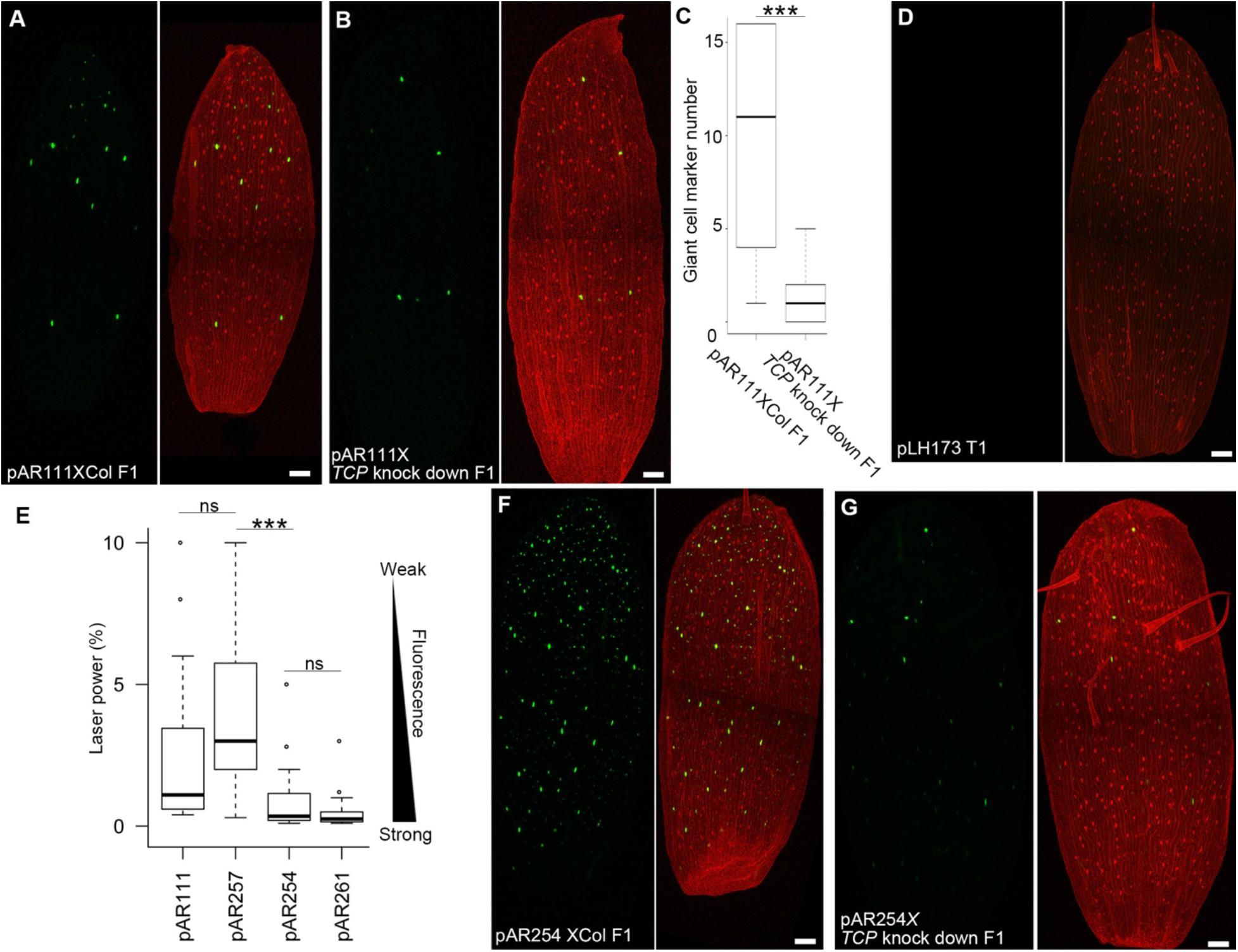
TCP transcription factors promote expression activity. **(A)** Confocal images of a sepal from *pAR111* and Col-0 (Col) hybrid plants. **(B)** Confocal images of a sepal from *pAR111* and *TCP* knock-down (*jaw-1D*) hybrid plants. **(C)** The number of nuclei expressing 3×Venus in *pAR111* × Col hybrid sepals and *pAR111*X *TCP* knockdown (*jaw-1D*) hybrid sepals. The boxes extend from the lower to upper quartile values of the data, with a line at the median. The whiskers extend past 1.5 of the interquartile range. ***p < 0.001, significant difference by two tailed t-test, p = 9.86E-08; n = 20. **(D)** Confocal images of a sepal from a *pLH173* (all TGGG putative TCP sites mutated in giant cell enhancer fragment 2) T1 plant. Hardly any nuclei expressing 3×Venus can be detected. **(E)** The laser intensity required to saturate the 3×Venus signal in giant cell for each reporter line. *pAR257* needs much higher laser intensity, indicating it has much lower expression level of 3×Venus. The boxes extend from the lower to upper quartile values of the data, with a line at the median. The whiskers extend past 1.5 of the interquartile range. ns, non-significant difference (t test); ***p < 0.001, significant difference by two tailed t-test, p = 1.07E-11; n = 17 for pAR111, n = 22 for pAR257, n = 24 for pAR254, and n = 24 for pAR261. **(F)** Confocal images of a sepal from a hemizygous *pAR254* plant (*pAR254* × Col F1). **(G)** Confocal images of a sepal from *pAR254 × TCP* knock-down (*jaw-1D*) F1 hybrid plants. Sepals were imaged at the same setting, with 3% laser power. Scale bars: 100 μm. See also Figures S3, S4, and S5.

DNA affinity purification sequencing (DAP-seq) analysis (O’Malley et al., 2016) indicated that TCP transcription factors bind to Region 2 (Figure S3) and throughout the intergenic region upstream of *AT5G17700*. However, there are no obvious canonical TCP binding sites in Region 2. We selected three sites with TGGG, which is half of the consensus TCP binding sequence, as putative TCP binding sites in Region 2 (Figure S4). When all three of the TGGG sites in the Region 2 reporter *pAR254* were mutated to TAAA (*pLH173*), the reporter lines expressed almost no Venus signal in sepal cells (Figure 3D and S5A). However, with extremely strong laser power, dim signal could be detected in some *pLH173* sepal cells (Figure S5B). This indicated that disruption of all the TGGG sites significantly reduced transcriptional activity in Region 2 (Figure 3D). Thus, our results suggest that binding of TCPs to Region 2 is critical for this region to activate expression.

### Lowering the expression level increased giant cell specificity

While we were dissecting the enhancer into regions (Figures 1-2), we noticed that different constructs required different laser intensity to generate the same fluorescence intensity of the reporter in giant cells (as determined by giant cell signal saturation). We recorded the laser intensity that different constructs required in order to reach the same giant cell fluorescence intensity and found that *pAR257* (Region 1-2) needed much higher laser intensity than *pAR254* (Region 2) alone to saturate the fluorescence in giant cells (Figure 3E), suggesting that Region 1 reduces transcriptional activity of Region 2.

Our analysis suggests that Region 1 reduces the activity of Region 2 and also limits reporter expression mainly to giant cells. We hypothesized that these two functions of Region 1 are connected such that globally reducing expression activity would be sufficient to shift a broad epidermal expression pattern toward a giant cell-specific expression pattern. To test this hypothesis, we lowered the transcriptional activity of Region 2 by crossing the Region 2 reporter (*pAR254*) to the *jaw-1D* line. Plants heterozygous for both *pAR254* and *jaw-1D* (F1) had sepals that displayed a giant cell-specific expression pattern (Figure 3G), instead of the broad epidermal expression pattern in sepals heterozygous for *pAR254* alone (Figure 3F; *pAR254* × wild-type F1 heterozygote), which confirmed that reducing Region 2 transcriptional activity increased the specificity of the expression pattern to giant cells. These data are consistent with the hypothesis that Region 1 limits the transcriptional activity driven by Region 2 to giant cells in part by reducing the overall expression level, thus reducing the relatively low expression in small cells (Figure 1G) and allowing only the relatively high expression in giant cells to persist.

### Dof transcription factors repress expression of the enhancer by lowering the expression level and limiting it to giant cells

We next searched for transcription factor binding sites in Region 1 responsible for repressing expression, which give rise to giant cell specificity. Our yeast one-hybrid screen showed that several Dof transcription factors bind to Region 1 and the overlap but not Region 2. Based on the AthaMap website (http://www.athamap.de) transcription factor binding site prediction, a 24 bp region 14 bp before the end of Region 1 contained three Dof transcription factor binding sites (AAAG) (Noguero et al., 2013; Sani et al., 2018). We generated a reporter construct containing Region 2 and extending 40 bp into the end of Region 1, which contained all three potential Dof binding sites (Region 2+Dof in *pAR307*, Figure 4A and S4). As a control, we also generated a construct containing Region 2 and 17 bp at the end of Region 1, which excluded the Dof binding sites (*pAR308*, Figure 4A and S4). The expression pattern of *pAR307* also varied among transgenic lines. Categorizing the expression pattern showed that *pAR307* retained most of the giant cell specific expression that Region 1-2 *pAR257* has (Figure 4B). Removing the Dof binding sites region led to an obvious shift in the expression pattern, with a majority of the *pAR308* plants showing epidermal expression (Figure 4B), indicating the importance of the Dof binding sites in generating the giant cell-specific expression pattern. Site-directed mutagenesis at these three putative Dof binding sites in the 40 bp region (Region2-Dof mutant, *pLH166*) reproduced the epidermal expression pattern in *pAR308* (Figure 4B). There is also a trihelix site in the 24 bp Dof binding site region, and mutation of the trihelix site did not alter the expression pattern (data not shown). We investigated the transcriptional activity of the enhancer fragments in *pAR307, pAR308*, and *pLH166*, by quantifying the laser intensity required for fluorescence to reach saturation in giant cells. The shift of giant cell specific expression to epidermal expression was correlated with an increase of reporter signal strength (detected with a lower power laser; Figure 4C), consistent with our previous observations (Figure 3E).

**Figure 4.**
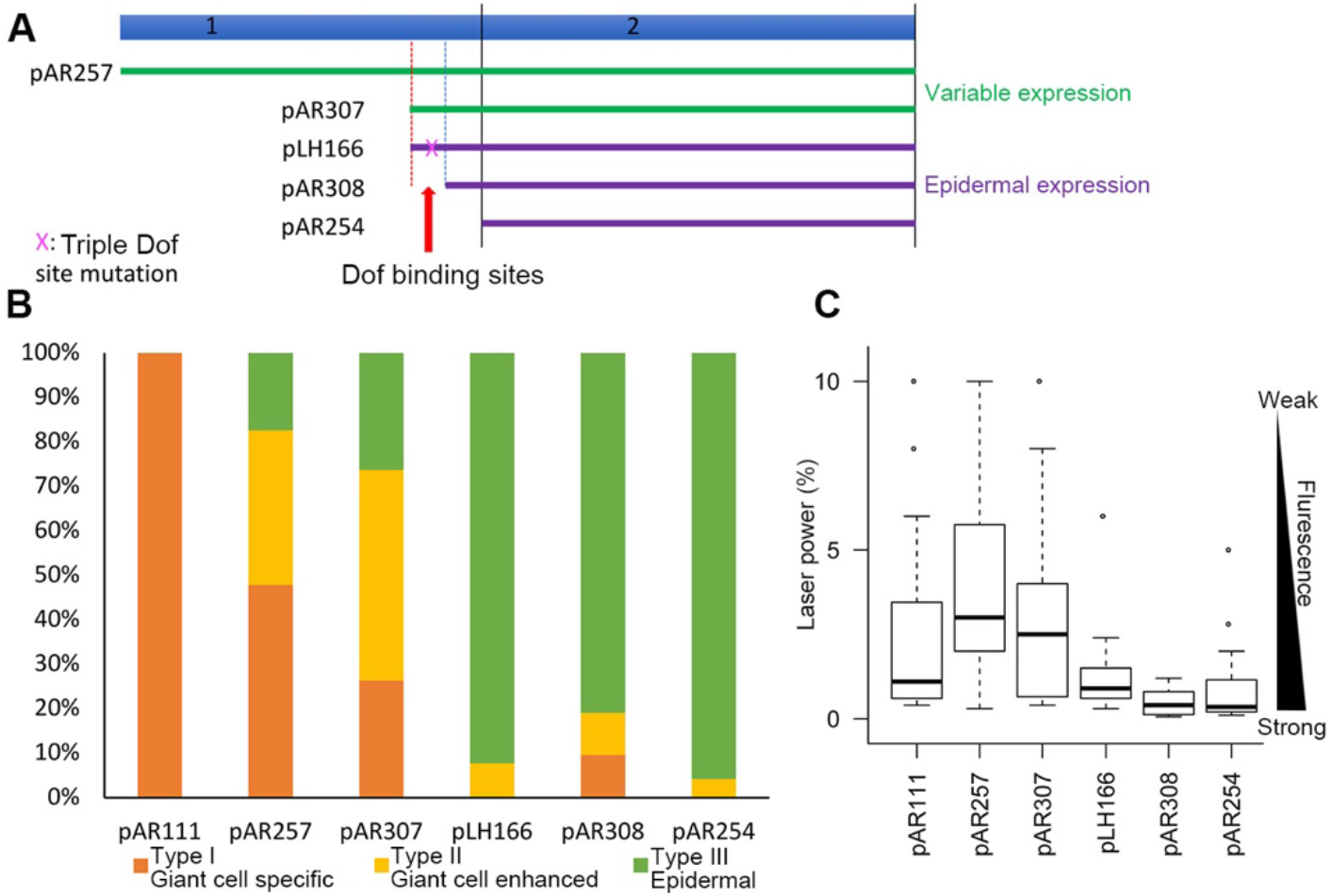
Dof binding sites in Region 1 confer giant cell enhanced activity. **(A)** Fine-scale dissecting of Region 1 in the enhancer. Schematics of reporter lines containing different fragments or mutation of Region 1 were shown. **(B)** Percentages of the three expression pattern types in different reporter lines. Data for *pAR111, pAR254* and *pAR257* were duplicated from Figure 2D for comparison. **(C)** The laser intensity required to saturate the 3×Venus signal in giant cell for each reporter line. The higher laser intensity signifies weaker fluorescence of the reporter. The boxes extend from the lower to upper quartile values of the data, with a line at the median. The whiskers extend past 1.5 of the interquartile range. Data for *pAR111, pAR254* and *pAR257* were duplicated from Figure 3E for comparison. See also Figure S4, S6, and S7.

Based on DAP-seq analysis, several Dof transcription factors bind to the 24 bp Dof binding site in Region 1 (Figure S6). The ones showing the highest binding affinity include: *AtDOF2.2* (*AT2G28810*), *HIGH CAMBIAL ACTIVITY2* (*HCA2 AT5G62940*), and *AtDOF5.8* (*AT5G66940*). We overexpressed each of the genes encoding these Dofs under the constitutive 35S promoter in the *pAR111* full length giant cell enhancer background to assess the impact of Dofs on enhancer activity. Overexpression of each of these Dofs strongly decreased *pAR111* expression in sepal giant cells (Figure 5D-L) compared with the sepal giant cells from the *pAR111* control (Figure 5A-C), indicating that Dof transcription factors repressed transcriptional activity of the giant cell enhancer. Overexpression of the Dofs also severely inhibited sepal development, resulting in shorter, narrow sepals in *35S::AtDOF2.2* and *35S::HCA2* (Figure 5D-E and G-H), and longer, narrow sepals in *35S::AtDOF5.8* (Figure 5J-K). Even in these morphologically altered sepals, giant cells were present, based on size and morphology. A couple of these giant cells expressed low levels of the reporter, indicating that repression of *pAR111* expression was not due to the absence of giant cells, but instead to repression of expression (arrowheads in Figure 5F, I, and L). Thus, our results demonstrate that binding of Dofs to Region 1 reduce overall reporter expression to confer giant cell-specific expression (Figure 4 and 5).

**Figure 5.**
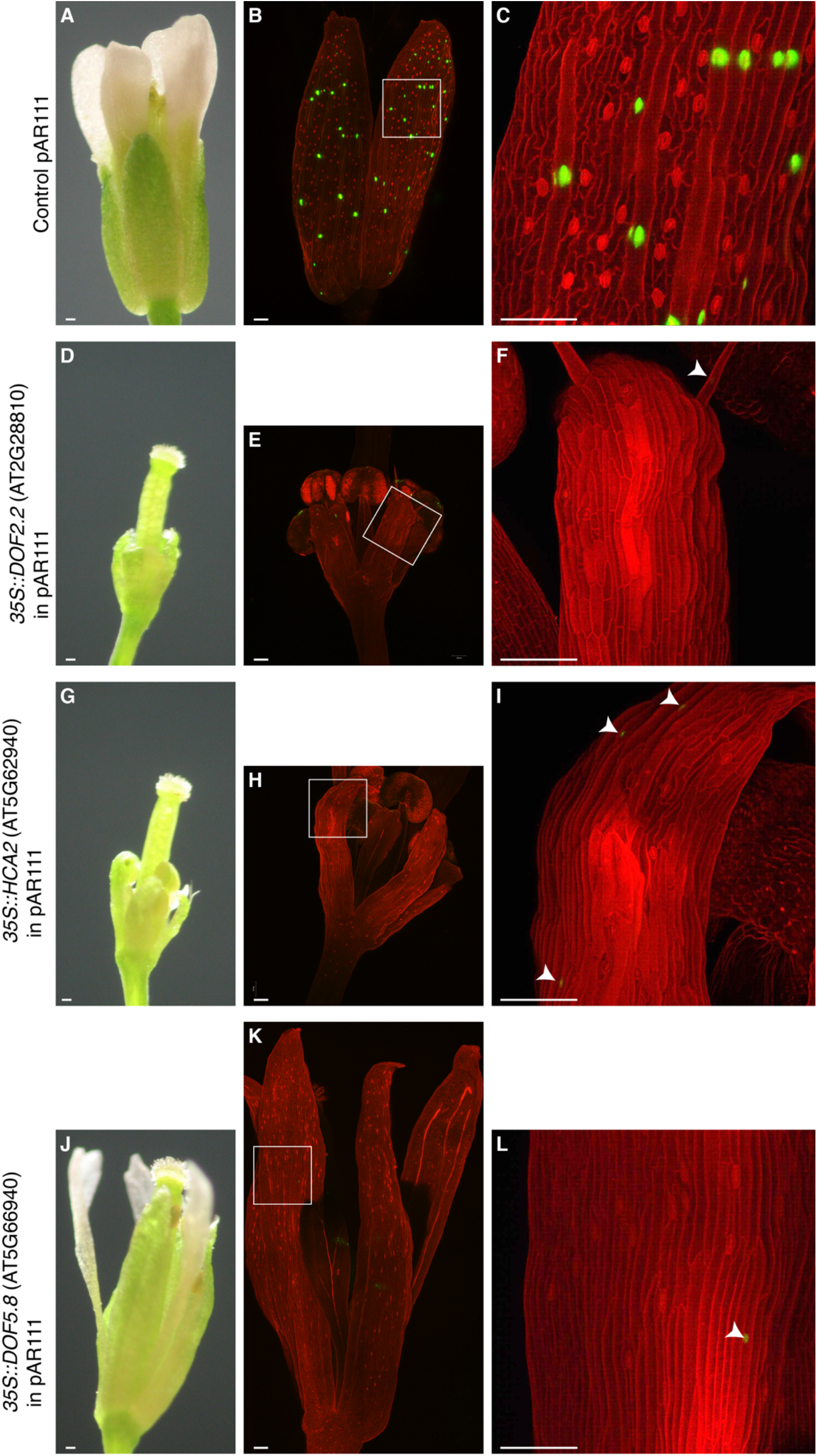
Dof transcription factor overexpression suppresses giant cell enhancer activity. *pAR111* is the full length giant cell enhancer which drives 3×Venus-N7 expression in giant cells (nuclear localized green signal). Cell walls were stained with PI (red). Arrowheads indicate nuclei expressing weak signal. **(A-C)** Flowers from a control *pAR111* plant. (A) A dissecting microscope image. (B) A confocal image showing bright *pAR111* full length giant cell enhancer reporter signal in giant cell nuclei. (C) Magnification of the white box in B. **(D-F)** Flowers from a *pAR111* plant expressing *p35S::DOF2.2* (*AT2G28810*). (D) A dissecting microscope image showing stunted sepals, petals, and stamens. (E) A confocal microscope image showing strong repression of the *pAR111* giant cell enhancer activity. (F) Magnification of the white box in E. **(G-I)** Flowers from *pAR111* plant expressing *p35S::HCA2* (*AT5G62940*). (G) A dissecting microscope image showing stunted sepals, petals, and stamens. (H) A confocal microscope image showing strong repression of the *pAR111* giant cell enhancer activity. (I) Magnification of the white box in H. **(J-L)** Flowers from *pAR111* plant expressing *p35S::DOF5.8* (*AT5G66940*). (J) A dissecting microscope image showing the sepals are narrow and highly elongated. (K) A confocal microscope image showing strong repression of the *pAR111* giant cell enhancer activity. (L) Magnification of the white box in K. Note that the arrowhead points to expression in a giant cell and that faint transverse walls that appear are in an underlying cell layer. Scale bars: 100 μm. Confocal images were taken with the same settings at 3% laser power. C, F, I, and L were rendered in MorphoGraphX adjusted in Photoshop with auto contrast. See also Figure S6.

## Discussion

In this study, we used giant cells as a model system to investigate how a cell type-specific expression pattern is achieved and reported the dissection of an enhancer region driving a giant-cell-specific expression pattern. This enhancer has a modular organization: Region 2 activated by TCP transcription factors drives expression in all epidermal cells with higher levels of expression in giant cells than small cells, while Region 1 is bound by Dof transcription factors to represses expression, thus limiting expression to giant cells (Figure 6). This repression contributes to cell type specificity due to the differences in expression level between giant cells and small cells driven by Region 2, which allows only the giant cell expression to remain after repression. Intriguingly, neither the activation nor the repression appears to be cell-type specific, yet the combination results in giant cell specificity.

**Figure 6.**
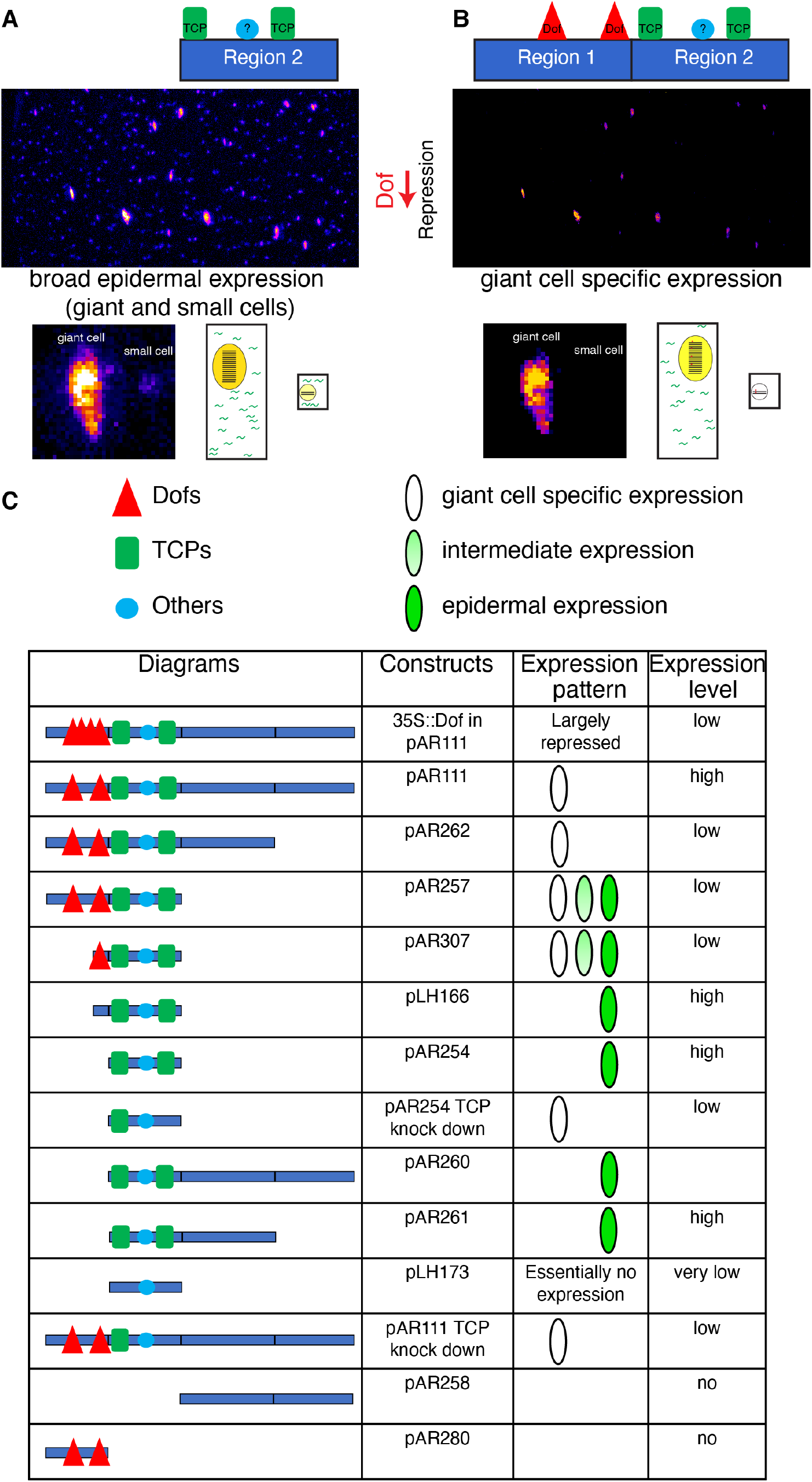
Summary: Giant cell specificity arises from the repression of a broad epidermal expression pattern. In the giant cell enhancer, cell type specificity of expression arises primarily from the combination of two elements: Region 1, an element driving broad epidermal expression, with higher expression levels in giant cells than small cells, and Region 2, an element repressing expression, and making the expression pattern giant cell specific. **(A)** Region 2 of the giant cell enhancer drives broad epidermal expression of the reporter. Nevertheless this expression is stronger in giant cells (large, highly endoreduplicated nuclei) than small cells (smaller nuclei) as highlighted by the fire LUT (using FIJI) applied in A. Our evidence suggests that TCP transcription factors (represented as green rectangles) bind to and activate expression via Region 2 of the enhancer. Note that our representation of two green rectangles representing TCPs, is meant to represent a wild type level of binding to the three putative TCP sites we identified, not a precise number of transcription factors bound. Our yeast-one hybrid results suggest that other factors (represented generally as a blue oblong) including HD-ZIP Class IV transcription factors bind to Region 2. We speculate these may be important in either activating epidermal expression or activating stronger expression in giant cells than small cells. One factor that may contribute to the increased expression in giant cells is the increased ploidy of these cells, providing many more copies of the enhancer (as represented in the giant cell with 16C DNA content) and lots of transcripts (represented in green) to drive strong enhancer expression (strong yellow in the nucleus) in comparison to the small diploid cell with few transcripts and low expression in the nucleus. **(B)** When combined with Region 1, Region 2 becomes much more giant cell specific as detailed in the results. Our evidence suggests that Dof transcription factors (represented as red triangles) bind to this element and repress expression. Note again, that the number of triangles is meant to convey the wild type occupancy of Dofs bound to Region 1, not an exact number of proteins. **(C)** Table summarizing the results of each of our dissection constructs together with site specific mutations and genetic manipulation of TCPs and Dofs. Again, the red triangles, green rectangles, and blue ovals represent relative occupancy, not exact numbers of transcription factors bound to DNA.

Rough dissection (Figure 6C) indicated that transcription factors interacting with Region 2 are candidates for activating broad epidermal expression. The results from the large-scale yeast one-hybrid assay suggest 37 transcription factors showed high-confidence interaction with this region, including proteins from the TCP gene family (Table 1). The TCP family is divided into two main classes, based on differences in the TCP domains and binding sites. In general, class II TCP proteins are attributed to functions that inhibit plant cell division and growth, whereas class I proteins promote proliferation. For example, class II protein CINCINNATA (CIN) controls leaf surface curvature in *Antirrhinum* by making cells more sensitive to an arrest signal, inhibiting proliferation (Crawford et al., 2004; Nath et al., 2003). Interestingly, class II protein TCP4 regulates both density and branching of trichomes, another highly endoreduplicated cell type in Arabidopsis (Vadde et al., 2017; 2019). Class I TCP15 regulates expression of boundary-specific genes (Uberti-Manassero et al., 2011) and TCP20 promotes cell division (Li et al., 2005). Intriguingly, TCP15 also regulates endoreduplication in Arabidopsis (Zi-Yu et al., 2011). Moreover, specific repression of TCP15 by fusing to an EAR repressor domain has been reported to result in a loss of giant cells in the abaxial sepal epidermis (Uberti-Manassero et al., 2011). The TCP proteins identified from the screen, which bound to Region 2, (TCP2, TCP3, and TCP4) are all in the class II CINCINATA subfamily (Martín-Trillo and Cubas, 2010). As proteins in this subfamily act to repress cell proliferation, especially at the leaf margin, it is plausible, though untested, that these factors serve to coordinate activation of expression at the enhancer with the transition from proliferation to differentiation, since the giant-cell enhancer is only active after cells have stopped dividing (Roeder et al., 2012).

Among the yeast one-hybrid candidates, another group of transcription factors that might be involved in giant cell specific expression were the three HD-ZIP Class IVs: HDG11, ANL2, and GL2. HD-ZIP Class IV transcription factors are typically restricted to the L1 and act to specify epidermal cell type identity (Nakamura et al., 2006), making these strong candidates for activating epidermal expression at the enhancer. HDG11 has been shown to have a small role in promoting giant cell formation (Roeder et al., 2012). ATML1, a HD-ZIP Class IV not included in the yeast one-hybrid assay, has been shown to be required for giant cell patterning as well as specification of epidermal identity (Abe et al., 2003; Meyer et al., 2017; Roeder et al., 2012). We have previously shown that overexpression of ATML1 is sufficient to create ectopic giant cells covering the sepal and that these ectopic giant cells express the *pAR111* full length giant cell enhancer reporter (Meyer et al., 2017), suggesting ATML1 may be involved in activating enhancer activity. Enhancer activity is downregulated but not completely absent in *atml1* mutants, consistent with a role for ATML1 in activation of enhancer activity (Roeder et al., 2012). The canonical binding site of ATML1 and other HD-ZIP Class IV transcription factors, the L1 box (TAAATGCA) (Abe et al., 2001; Nakamura et al., 2006), does not appear in the enhancer sequence. However, we speculate that HD-ZIP Class IV proteins may recognize another motif in the enhancer or interact with other transcription factors. Non-L1-box regions have been shown to specify epidermal identity; the promoter of ATML1 activates epidermis-specific expression even when its L1 box is deleted (Takada and Jurgens, 2007).

It is currently unclear what gives rise to higher expression in giant cells than small cells in the broad epidermal expression pattern generated by Region 2. Higher expression in giant cells could be caused by a combination of factors. One is that giant cells are endoreduplicated, generally reaching a ploidy of 16C; thus, higher DNA content and associated increases in cell size could enhance expression (Robinson et al., 2018; Roeder et al., 2010). In trichomes, endoreduplication has been shown to be important for the maintenance of cell identity (Bramsiepe et al., 2010). However, as we have previously described, this giant cell enhancer is not strictly a marker of ploidy and additional factors contributing to increased expression in giant cells are likely (Roeder et al., 2012). HD-ZIP Class IV transcription factor ATML1 reaches higher levels of expression in giant cells than small cells during their specification, which could initiate this pattern of higher expression (Meyer et al., 2017). Finally, it is possible that TCPs or other factors binding to Region 2 generate this difference in expression level. Further elucidating the cause of this difference in expression will be an important future goal in understanding the relationship between endoreduplication and cell identity in giant cells.

Giant cell specificity arises through the repressive activity of Region 1 (Figure 6C). Both yeast one hybrid and DAP-seq results suggest that Dof transcription factors bind to Region 1 and overexpression analyses and enhancer dissections suggest Dofs repress transcription to create giant cell specificity. Region 1 alone does not activate any expression, suggesting it acts solely as a repressor and detecting its output requires the activators bound to Region 2. Dof transcription factors bind AAAG sites in DNA through a single N-terminal C_2_C_2_-zinc-finger like motif (Noguero et al., 2013). Dof proteins interact more tightly with the DNA when two proximate binding sites are present than they do with only one binding site (Sani et al., 2018), which is consistent with the three nearby putative Dof sites in Region 1. There are 36 Dof family members in *Arabidopsis* which tend to function redundantly (Gupta et al., 2015; Noguero et al., 2013; Yanagisawa, 2002; 2015). Overexpression of each of the three *Dof* genes we tested was sufficient to repress expression of the giant cell enhancer. Dof overexpression also altered sepal development, making sepals narrow, and in most cases shorter, suggesting that these *Dof* genes are important for sepal morphogenesis, not just regulation of this enhancer. Dofs have been shown to regulate many developmental processes and environmental responses including seed germination, light response, vascular cambium development, and cell division (Noguero et al., 2013). Dofs can act as either activators or repressors in different contexts (Noguero et al., 2013). The Dof transcription factor OPB1 promotes cell cycle entry by directly activating CYCLIN D3;3 and AtDOF2.3 which is associated with replication (Skirycz et al., 2008). In seeds, DOF3.2 binds to the DELLA co-repressor RGA-like 2 (RGL2) and DOF AFFECTING GERMINATION 1 (DAG1) binds to the DELLA corepressor GA INSENSITIVE (GAI) to repress seed germination in the absence of gibberellin (GA), linking Dofs with GA signaling (Boccaccini et al., 2014; Ravindran et al., 2017; Ruta et al., 2020). This observation raises the possibility that the repressive activity conferred by the Dofs could be mediated by co-repressors. Of the Dofs we tested, the functions of DOF2.2 (AT2G28810) remain unknown (Yanagisawa, 2015). HCA2/DOF5.6 (AT5G62940) promotes the development of the vascular cambium and consequently the radial growth of the root (Guo et al., 2009; Miyashima et al., 2019). DOF5.8 (AT5G66940) regulates ANAC069 in response to abiotic stress (He et al., 2020). In the future, it will be interesting to determine which Dofs are required for the repressive activity of Region 1 in wild type sepals, since our overexpression analysis have shown that all of three Dofs we tested, despite coming from different clades, were sufficient to repress activity (Yanagisawa, 2002; 2015). Given the binding of multiple Dofs, including the three we tested, to this enhancer region in the DAP-seq database, we expect multiple Dof proteins to act redundantly in this role.

Enhancers tend to be conserved across closely related species because selection preserves their function whereas nonfunctional intergenic sequences are not under selection and diverge faster. This conservation has been used to identify enhancer sequences in non-coding DNA (Pennacchio et al., 2006). The giant cell enhancer includes a previously identified conserved noncoding sequence (Haudry et al., 2013). Alignment of Region 1-2 enhancer sequences from six species across the Brassicaceae family revealed that two of the three putative Dof binding sites we identified are conserved, and all three of the putative TCP binding sites are conserved in *Arabidopsis lyrata, Cardamine hirsuta, Brassica rapa, Sisimbrium irio*, and *Eutrema salsugineum* (Figure S7). In the future, it will be interesting to determine whether these enhancers produce similar expression patterns in these other species.

Previous studies in animals have often shown spatial restriction of enhancer activity is generated by broad activation combined with localized repression (Spitz and Furlong, 2012). For example, during early stages of sea urchin development, the *endo16* gene is precisely expressed in the endoderm through broad activation in the embryo by the enhancer Region A combined with localized repression in the neighboring ectoderm by Regions E and F and localized repression in the neighboring skeletogenic mesenchyme by Region DC (Davidson, 2010). Likewise, in *Drosophila* the *even-skipped* (*eve*) stripe two enhancer is activated broadly by Bicoid (Bcd) and Hunchback (Hb), and narrowed to a precise stripe through repression by Giant (Gt) on the anterior and Krüppel (Kr) on the posterior (Stanojevic et al., 1991). Genetic analysis suggests a similar regulatory logic is found in Arabidopsis during the establishment of the narrow band of *SHATTERPROOF* (*SHP*) expression precisely at the valve margin through repression by FRUITFULL (FUL) in valves and REPLUMLESS (RPL) in the replum (Ferrándiz et al., 2000; Roeder et al., 2003). Our findings in the giant cell enhancer are distinct in that both the activator and repressor do not appear to be spatially specific, and yet the combination gives rise to cell type specificity. Broad activation of the enhancer in all epidermal cells through a combination of TCP and other transcription factors likely including HD-ZIP Class IV transcription factors generates a pattern with high expression in giant cells and low expression in small cells. The Dof transcription factors repress expression by lowering levels until the enhancer no longer activates expression in small cells, and only expression in giant cells remains detectable (Figure 6). Thus, we suggest that cell-type specificity can emerge from the combination of two modules neither of which is itself cell-type specific.

## Methods

### Plant Growth

*Arabidopsis thaliana* Col-0 plants were grown on Lambert Mix LM-111 soil in Percival growth chambers at 22 °C with 24 hours fluorescent light (~100 μmol m^−2^ s^−1^) conditions. Seeds were planted on soil and cold stratified for about two days before placing in the growth chambers.

### Giant cell enhancer dissection

Cloning of the full length 1024 bp enhancer construct pAR111 and verification that it was sufficient to drive giant cell-specific expression in either orientation was previously described (Roeder et al., 2012). This 1024 bp region was divided into four arbitrary regions with Region 1 falling closest to the downstream gene *AT5G17700* and farthest from the original enhancer trap insertion YJ158 (Figure 1). Enhancer fragments were tested for their activity in the orientation they would have relative to downstream gene *AT5G17700* such that Region 1 is closest to the reporter gene.

We first cloned a gateway destination reporter vector (pSL12) so that we could rapidly clone and assay the reporter expression patterns driven by fragments of the giant cell enhancer. pSL12 contains a gateway site upstream of the -60 minimal promoter form 35S and the 3×Venus-N7 super-bright yellow-fluorescent nuclear-localized reporter in the pMLBart binary vector backbone. pSL12 confers Basta resistance in plants and spectinomycin resistance in bacteria. To create pSL12, the gateway site in which the *Not*I restriction site had been removed was amplified with oAR505 and oXQ6 and cloned into pGEM-T Easy (Promega) to create pSL10. The gateway site was cut from pSL10 with *Xho*I and *Kpn*I and cloned into -60 3×Venus-N7-BJ36 (Roeder et al., 2012) to create pSL11. pSL11 was cut with *Not*I and the gateway -60 3×Venus-N7 fragment was cloned into pMLBart to create pSL12.

Fragments of the enhancer were amplified through PCR with Pfu Ultra II (Agilent) or Phusion (NEB) from the *Arabidopsis thaliana* genome BAC clone MAV3 or Col-0 genomic DNA using the primers specified in Table S2 for each fragment. The primers sequences are listed in Table S3. The PCR products were cloned into pENTR D TOPO (ThermoFisher) according to the manufacturer’s instructions to create the entry clones listed in Table S2. LR reactions (ThermoFisher) between the entry clones and pSL12 generated the final constructs listed in Table S2. Putative Dof and TCP sites were mutated through changes in the primer sequences. All constructs were verified by sequencing the inserts.

These constructs were transformed into Columbia (Col-0) plants through *Agrobacterium*-mediated floral dipping and were selected for Basta Resistance. For most constructs the expression patterns in approximately 20 T1 transgenic plants were analyzed. Note the original pAR111 transgenic plants described in (Roeder et al., 2012) were in the Landsberg *erecta* (L*er*) accession, but for this project pAR111 was transformed into Col-0. Transformation of the empty vector pSL12 into plants generated no expression (Figure S2D). As typical with the independent insertion of the T-DNA into the genome (Schubert et al., 2004), we saw differences in expression level and sometimes pattern between different T1 lines as characterized in the results. However, within a single plant the expression pattern was similar across sepals. Likewise the expression pattern was consistent in subsequent T2 and T3 generations for those lines examined.

### Confocal Microscopy

To minimize the morphological or gene expression variability caused by different plant vitality, the 12th to 25th flowers on the main stem were used for observation (Hong et al., 2016). Mature sepals (stage 14 according to (Smyth et al., 1990)) were dissected from the flowers with tweezers and needles and stained for 15 minutes in propidium iodide (PI; 0.15 mg/ml in water), mounted on slides in 0.01% triton X-100 under a cover slip, and imaged with a Zeiss 710 confocal laser scanning microscope. A 514 nm excitation laser was used to excite both the fluorophores. The 3×Venus-N7 enhancer reporter signal was collected between 519 to 566 nm, while the PI signal was collected between 603-650 nm. Images were taken using 10X (Plan-APOCROMAT NA = 0.45 air) or 20X (Plan-APOCROMAT NA = 1.0 water dipping) objective lenses. Different T1 transgenic lines have different levels of expression likely due to effects of the position of insertion of the T-DNA. For the experiments in Figures 1, 2, and 4 we varied the laser power to achieve saturation of the reporter in a few of the brightest giant cell nuclei (the other settings including gain were kept the same). This allowed us to compare reporter expression between cells while keeping the expression level in giant cells relatively constant (ranges of laser power used are reported in Figure 3E). In contrast, Figure S5 shows the same sepal imaged with different laser powers (3% and 40% as indicated in the figure), to detect very low levels of expression. For most images the maximum intensity projection is shown. For Figure 5C,F,I, and L the image was volume rendered in MorphoGraphX and subsequently Auto Contrast was used in photoshop to visualize the epidermal cell layer without the underlying layers.

### Quantitative Image Processing

FIJI (https://fiji.sc) with the Costanza plugin (http://home.thep.lu.se/~henrik/Costanza/) was used to quantify the fluorescence intensity and size of nuclei expressing pAR254. In the images used, the fluorescence intensity of the reporter in the nuclei was not saturated. The .lsm stack image from the microscope was opened in FIJI. The color of Channel 1 (with the reporter expression) was converted to greys using the Channels tool. The channels were split using Split Channels. The maximum intensity projection of Channel 1 (reporter expression) was created with Z project. The maximum intensity projection image was analyzed with the Costanza plugin with the following settings. In the general menu, “Mark intensity plateau with single maximum” was checked as were “Mark cell centers. Marker pixel radius: 3”, Display basins of attractions (BOA)”, and “Display basins of attractions according to measured intensity.” In the pre-processor queue, “Background extraction” with an intensity threshold of 20 was executed first followed by “Mean filter” with radius 0.1 and number of times 10. In the post-processor, “BOA remover” with a size threshold of 10 and an intensity threshold of 10 was executed first followed by a “BOA merger” with a radius of 15. Use ImageJ stack calibration was used for scaling. The results were analyzed in Microsoft Excel. Giant cells have enlarged nuclei, so a threshold of 100 μm^2^ was used to separate giant cell from small cell nuclei.

To count the nuclei number in Figure 3C, we kept the image settings including the laser power and gain the same when imaging the sepals for both *pAR111* × Col F1 and *pAR111 × jaw-1D* TCP knock down F1. Images were processed with maximal intensity projection using the Zeiss Zen software and nuclei which showed saturation of the reporter signals were counted as giant cell nuclei.

### Statistics

Two-tailed students T-tests were used for pairwise comparisons.

### High Throughput Yeast-One-Hybrid Screen

Yeast-one-hybrid screens were conducted with a nearly genome-wide transcription factor collection arrayed in 384 well plates (Pruneda-Paz et al., 2014) according to the protocol in (Li et al., 2019) using a luciferase reporter (Bonaldi et al., 2017). Bait plasmids were created through LR reactions of entry clones containing Region 1, Region 2 or a region overlapping the junction between Regions 1 and 2 into the pY1-gLUC59_GW yeast reporter plasmid (Table S4). The resulting constructs were integrated into the URA3 locus on the chromosome of the yeast strain YM4271.

### Genetic interaction with *jaw-1D*

The *jaw-1D* mutant (CS6948) was ordered from the Arabidopsis Biological Resource Center (ABRC). The homozygous *jaw-1D* mutant was crossed to *pAR111* plants and to *pAR254* plants. The expression patterns were analyzed in the F1 generation which were heterozygous for the reporter and the *jaw-1D* dominant mutant. The expression patterns were compared with the F1 generation of the cross of each reporter with Col-0 wild type as a control.

### Dof overexpression

Entry clones containing Dof family member cDNAs were ordered from the ABRC: TOPO-U09-G04 (AT2G28810), TOPO-U20-E12 (AT5G62940), and TOPO-U13-H01 (AT5G66940). Through LR reactions these were recombined into the binary 35S destination vector pK7WG2 to make pLH160 (35S::AT2G28810 CDS), pLH162 (35S::AT5G62940 CDS), and pLH163 (35S::AT5G66940 CDS). These constructs were transformed into pAR111 (in Col-0) plants through agrobacterium-mediated floral dipping and were selected for kanamycin resistance.

### Flower pictures

Images of flowers were taken on a Zeiss Stemi 2000-C stereomicroscope with a Cannon Powershot A640 digital camera.

### DAP-seq database analysis

The C_2_C_2_-Dof and TCP transcription factor families DAP-seq data was analyzed from the Plant Cistrome Database (http://neomorph.salk.edu/dev/pages/shhuang/dap_web/pages/index.php (O’Malley et al., 2016)) using the Genome Browser to examine the giant cell enhancer region on chromosome 5 coordinates 5837111-5837559 for Region 1-2.

### Alignment of the giant cell enhancer from Brassicaceae species

A conserved noncoding sequence (Chr5:5,837,330-5,837,417) has been identified among Brassicaceae species (Haudry et al., 2013) overlapping with Region 2 of the giant cell enhancer (http://mustang.biol.mcgill.ca:8885/cgi-bin/hgGateway). To examine sequence conservation of Regions 1 and 2 of the enhancer and the putative Dof and TCP binding sites we aligned the corresponding sequences from *Arabidopsis thaliana, Arabidopsis lyrata, Cardamine hirsuta* (CoGe id 36106 Cardamine hirsute v1 unmasked Chromosome 6 18239653-18239154), *Brassica rapa* (COGE id 24668 Brasica DB Chr unmasked v1.5 chromosome A10 11517566-11517500 and A02 3259372-3259603), *Eutrema parvulum* (formerly *Thellungiella parvula* 12384 UIUC unmasked v2 chomosome 6-6 5884609-5883951), *Eutrema salsugineum* (Id 19492 JGI unmasked 6007159-6007397), *Sisimbrium irio* (Id 20245 VEGI unmasked vVEGI 2.5 Chromosome scaffold_57 2159183-2159617). All sequences were checked that they fell in syntenic regions upstream of a homolog of *AT5G17700*. The sequences were aligned with Clustal Omega (https://www.ebi.ac.uk/Tools/msa/clustalo/) and the alignments were formatted and displayed with Boxshade (https://embnet.vital-it.ch/software/BOX_form.html). Alignments were adjusted slightly by hand around gaps and the ends of sequences were trimmed or extended to give the best alignment (Figure S7).

### Accession Numbers

Giant cell enhancer associated MATE efflux family protein, AT5G17700.

*jaw-1D* seed (CS6948) and associated gene *miR319a* (*AT4G23713*)

Dof family members analyzed: AT2G28810, AT5G62940, AT5G66940

### Distribution of Materials and Data

Requests for materials and data should be addressed to Adrienne H. K. Roeder.

## Supporting information

Dataset S1 Yeast one-hybrid for Region 1

Dataset S2 Yeast one-hybrid for Region 2

Dataset S3 Yeast one-hybrid for Region Overlapping the boundary between Region 1 and 2

## Author Contributions

Conception and design of experiments: L.H., C.S.K., A.H.K.R.

Giant cell enhancer dissection experiments and analysis: L.H., C.S.K., A.H.K.R.

High throughput yeast-one-hybrid screen: S.E.K., J.L.P.-P.

Writing of the manuscript: L.H., A.H.K.R.

Revising and editing of the manuscript: L.H., C.S.K., J.L.P.-P., A.H.K.R.

## Acknowledgements

We thank Joseph Cammarata, Frances Clark, Kate Harline, Shuyao Kong, Xiaobo Zhao and Ming Zhou for critical reading and comments on the manuscript. We thank Samuel Leiboff and Dana Robinson for technical assistance in cloning constructs for this project.

This work was supported by NSF IOS Plant, Fungal, and Microbial Developmental Mechanisms grants IOS-1256733 (AHKR) and IOS-1553030 (AKHR), NSF IOS 1755452 (JLPP), NSF MCB 1158254 (JLPP) and NIH R01GM056006 (JLPP). No conflicts of interest declared.

## Supplemental Information

Dataset S1: Yeast-one-hybrid results for Region 1.

Dataset S2: Yeast-one-hybrid results for Region 2.

Dataset S3: Yeast-one-hybrid results for the overlap between Regions 1 and 2.

**Figure S1.**
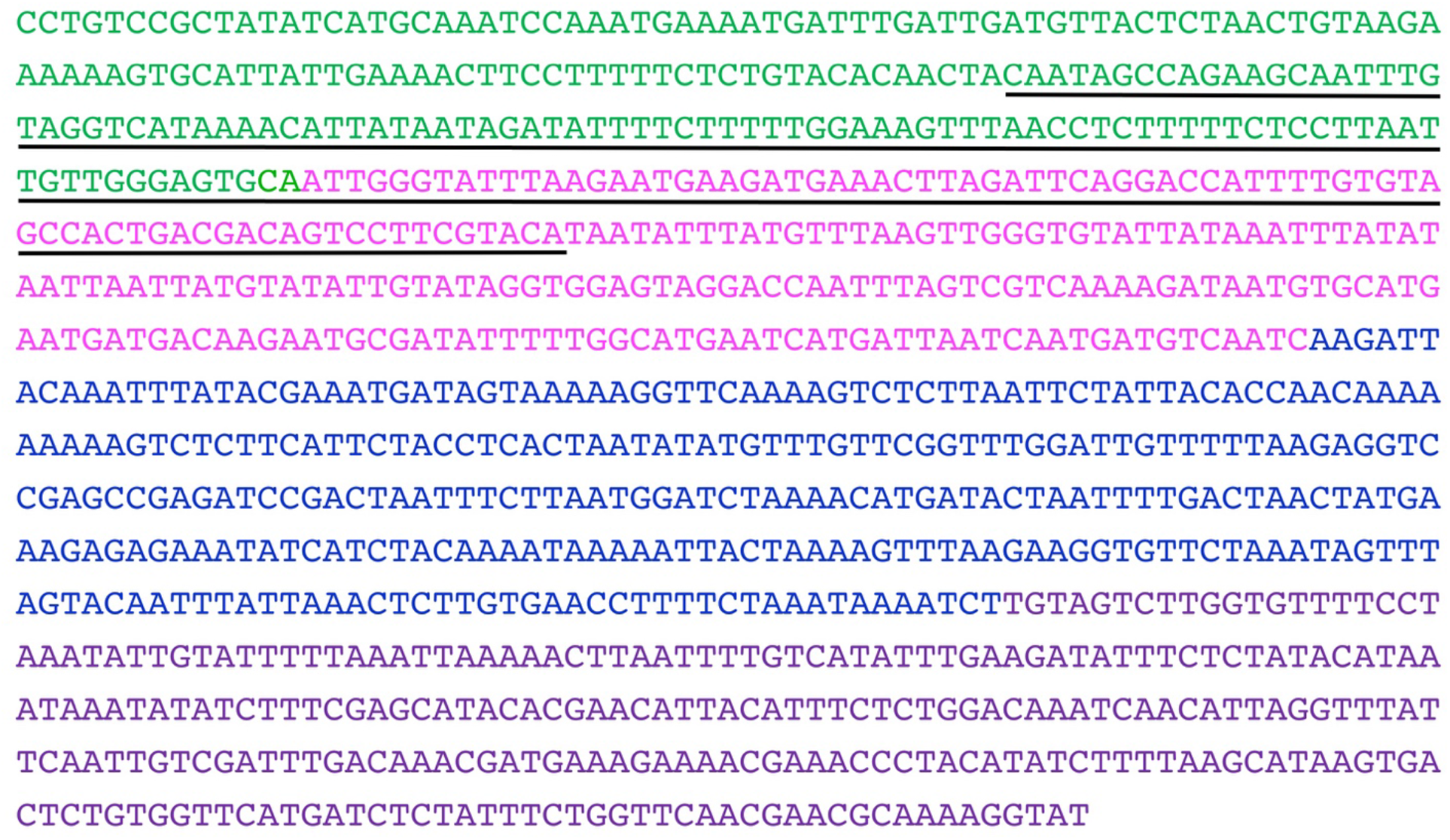
The whole sequence of the giant cell enhancer, related to Figure 1. Green: Region 1, 1-208 bp; magenta: Region 2, 209-450 bp; blue: Region 3, 451-760 bp; purple: Region 4, 761-1024 bp. The underlined region is the overlapping region used in the yeast one hybrid assay.

**Table S1.** Full-genome yeast one-hybrid screen results: low-confidence interactions, related to Table 1.

| Gene family | Region 1 | Overlap | Region 2 |
| --- | --- | --- | --- |
| **C<sub>2</sub>C<sub>2</sub>-Dof** | AT2G28810 | DOF4.5 (AT4G21080) |  |
| **TCP** |  |  | TCP13 (AT3G02150)<br>TCP22 (AT1G72010) |
| **AP2** | RAP2.4 (AT1G78080)<br>AT1G28160<br>AT4G39780 | AT5G60142 | CRF6 (AT3G61630)<br>DEWAX (AT5G61590) |
| **bHLH** |  |  | AKS2 (AT1G05805) |
| **others** | EIL3 (AT1G73730)<br>IAA26 (AT3G16500)<br>IAA31 (AT3G17600)<br>MYB27 (AT3G53200)<br>NAC038 (AT2G24430)<br>SPL2 (AT5G43270)<br>AT3G10030 | AGL14 (AT4G11880)<br>BZIP6 (AT2G22850)<br>POSF21 (AT2G31370)<br>SIG3 (AT3G53920)<br>SUVR2 (AT5G43990)<br>WOX11 (AT3G03660)<br>AT3G52250<br>AT4G00390<br>AT5G28040 | AGL55 (AT1G60920)<br>BBX10 (AT3G21880)<br>BPC7 (AT2G35550)<br>DAZ2 (AT4G35280)<br>EMB2746 (AT5G63420)<br>GPL1 (AT2G25650)<br>HDG5 (AT5G46880)<br>NAC078 (AT5G04410)<br>NAC098 (AT5G53950)<br>TGA2 (AT5G06950)<br>UNE16 (AT4G13640)<br>AT1G55760<br>AT1G60240<br>AT1G61730<br>AT1G66420<br>AT3G02400 |

**Figure S2.**
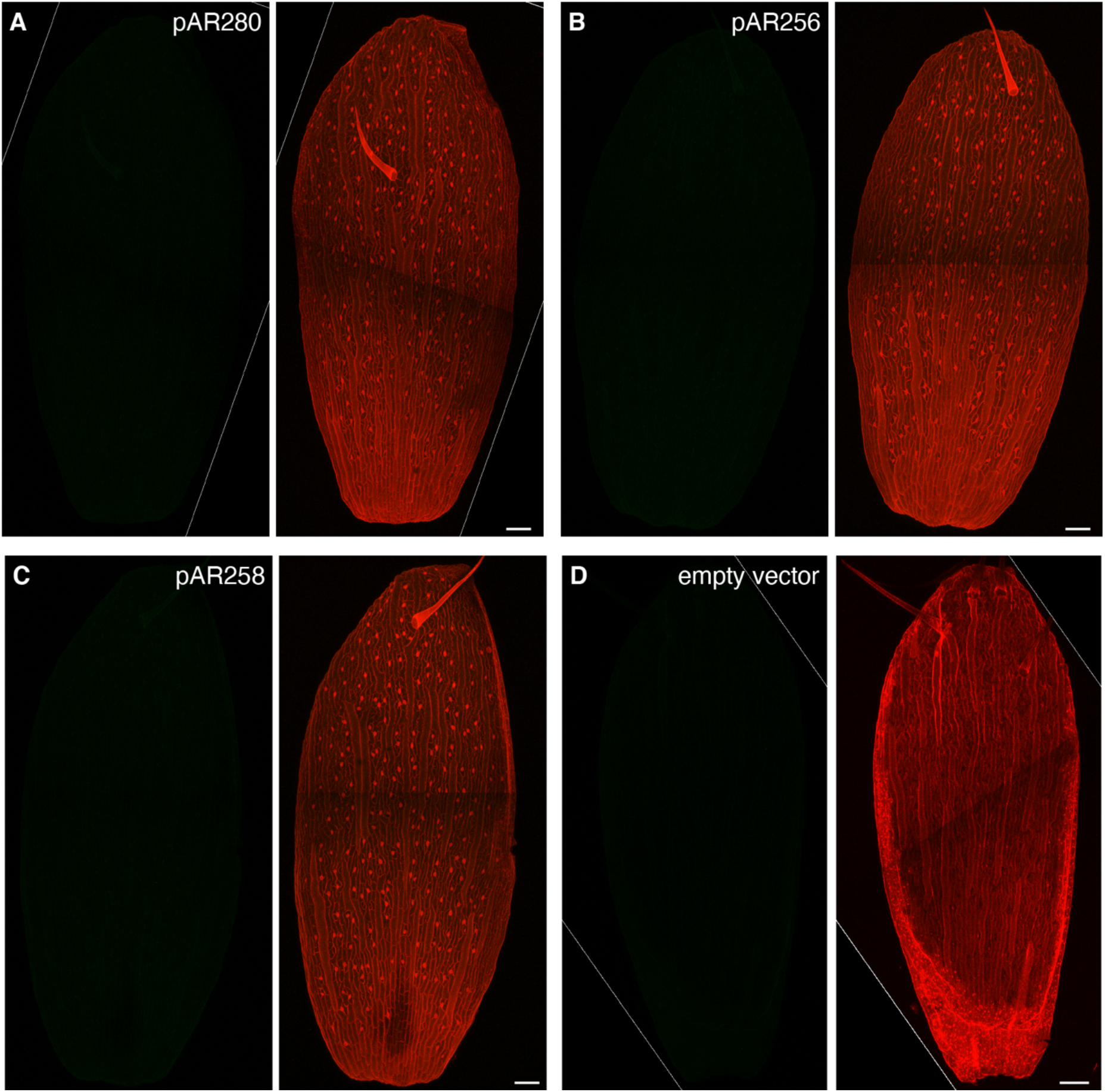
Region 1 alone, Region 4 alone and Regions 3 and 4 together are not sufficient to drive any reporter expression, related to Figure 1. Confocal images of stage 14 sepals from different reporter lines. Images on the left show 3×Venus (green) signal marking the nuclei of cells expressing the reporter. Images on the right show 3×Venus (green) signals merged with PI (red) stained cell walls. Images were taken with 3% laser power except where noted. Increasing laser power to 40% did not reveal any expression. **(A)** *pAR280* Region 1 alone does not drive any reporter expression. **(B)** *pAR256* Region 4 alone does not drive any reporter expression. **(C)** *pAR258* Region 3-4 do not drive any reporter expression. **(D)** The empty vector (*pSL12*) does not drive any reporter expression. Imaged at 2.6% laser power. Scale bars represent 100 μm.

**Fig. S3.**
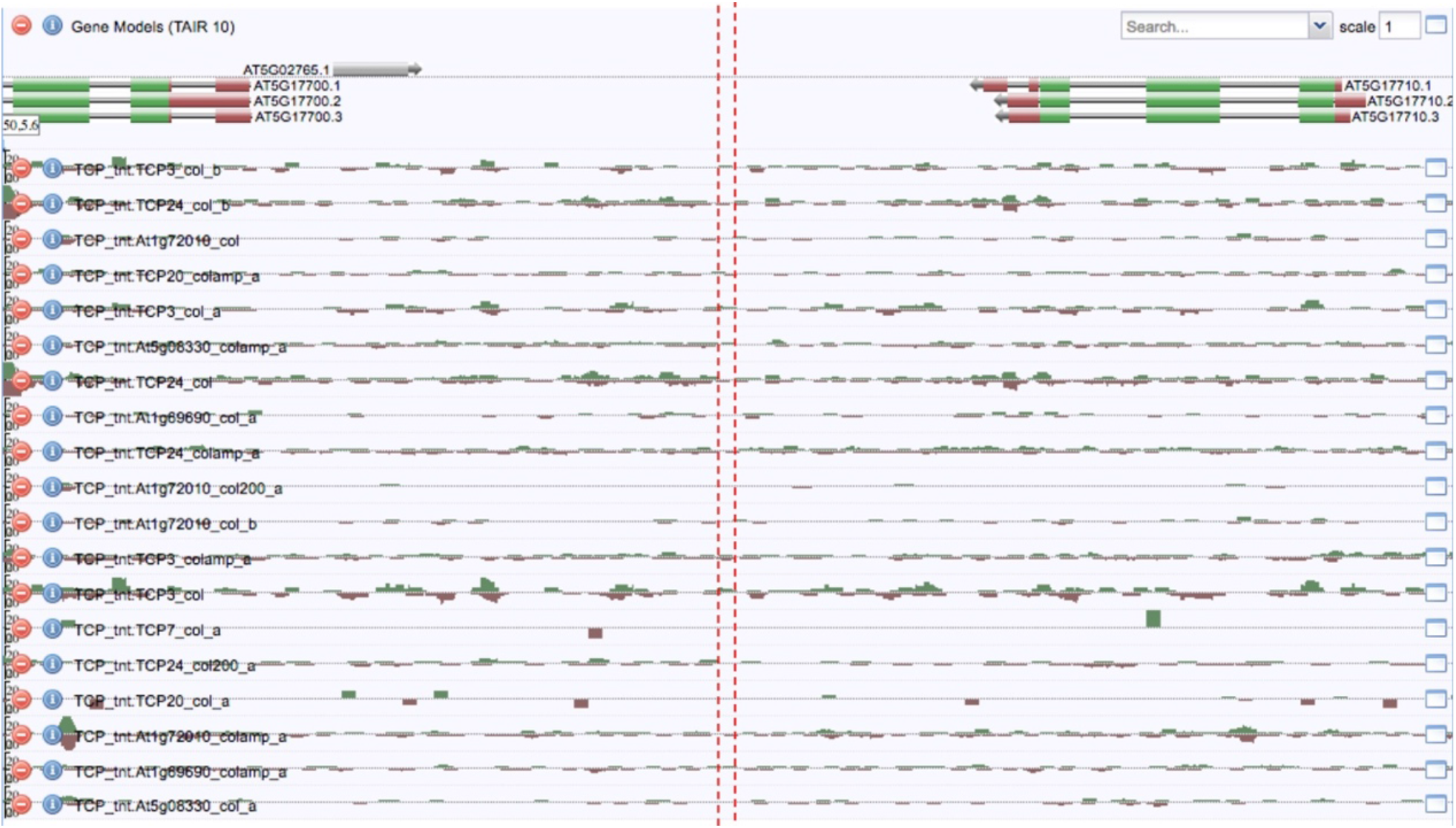
TCP transcription factors bind to the giant cell enhancer and nearby region, related to Figure 3. Binding sites of TCP transcription factors in the giant cell enhancer and nearby region. The red dash lines flank the three TCP binding site region in Region 2. Data are from DNA affinity purification sequencing (DAP-seq) analysis (http://neomorph.salk.edu/dev/pages/shhuang/dap_web/pages/aj_browsers.php). We observe moderate binding throughout the intergenic region including the region of the enhancer. We speculate that this moderate binding might reflect spatial restriction of binding either to sepals or epidermal cell layers, that would not be accurately captured by DAP-seq.

**Fig. S4.**
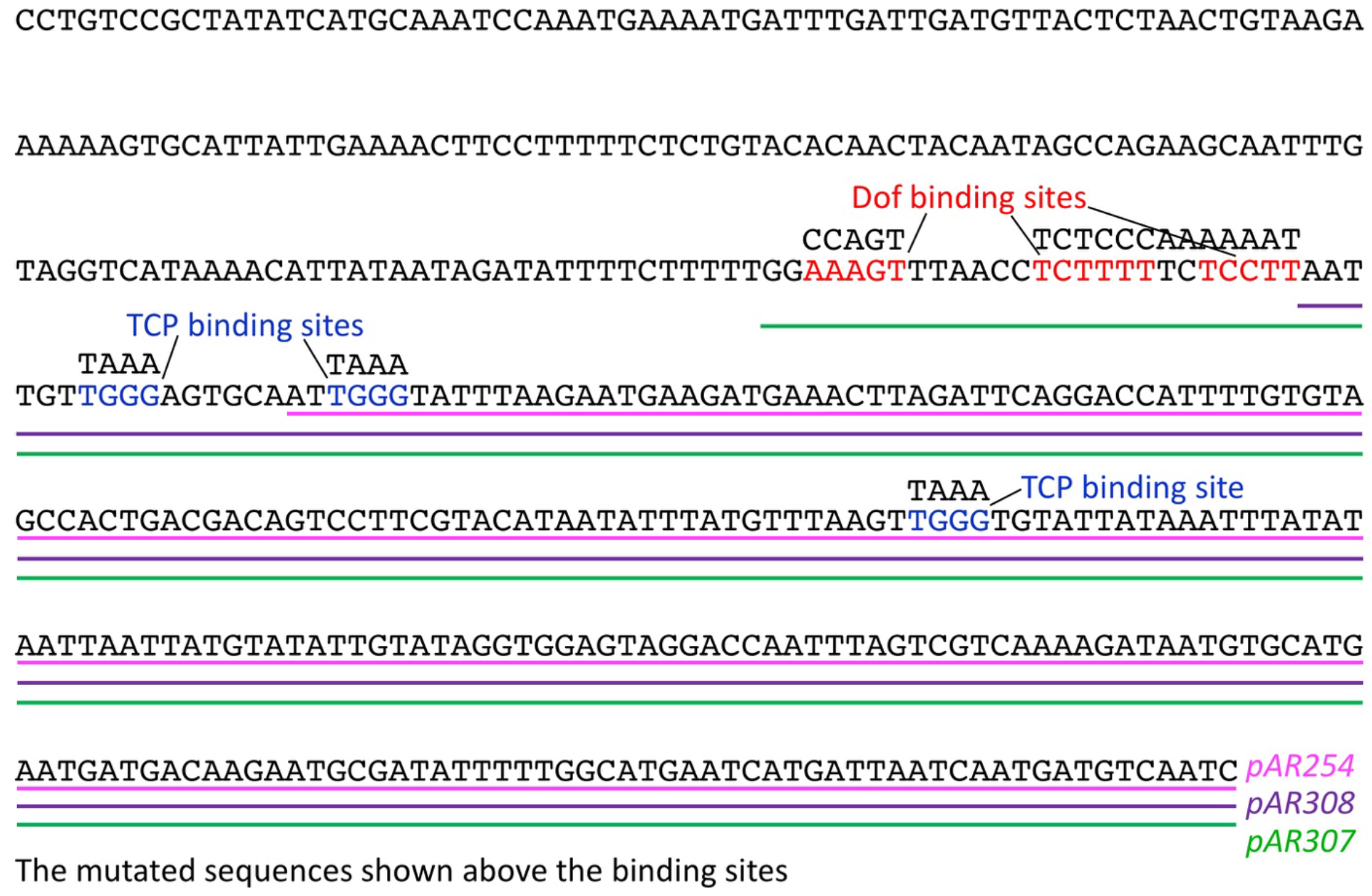
Sequences and locations of the putative Dof binding sites, putative TCP binding sites and different enhancer fragments, related to Figures 3 and 4. The sequences included in *pAR307* (Region 2 + Dof, green line), *pAR308* (Region 2 plus, purple line), and *pAR254* (Region 2 only, magenta line) are marked with lines below the sequence. The putative Dof binding sites are highlighted with red letters. The putative TCP binding sites are highlighted in blue. The mutated sequences for the putative Dof and TCP binding sites are shown above the original sequences.

**Figure S5.**
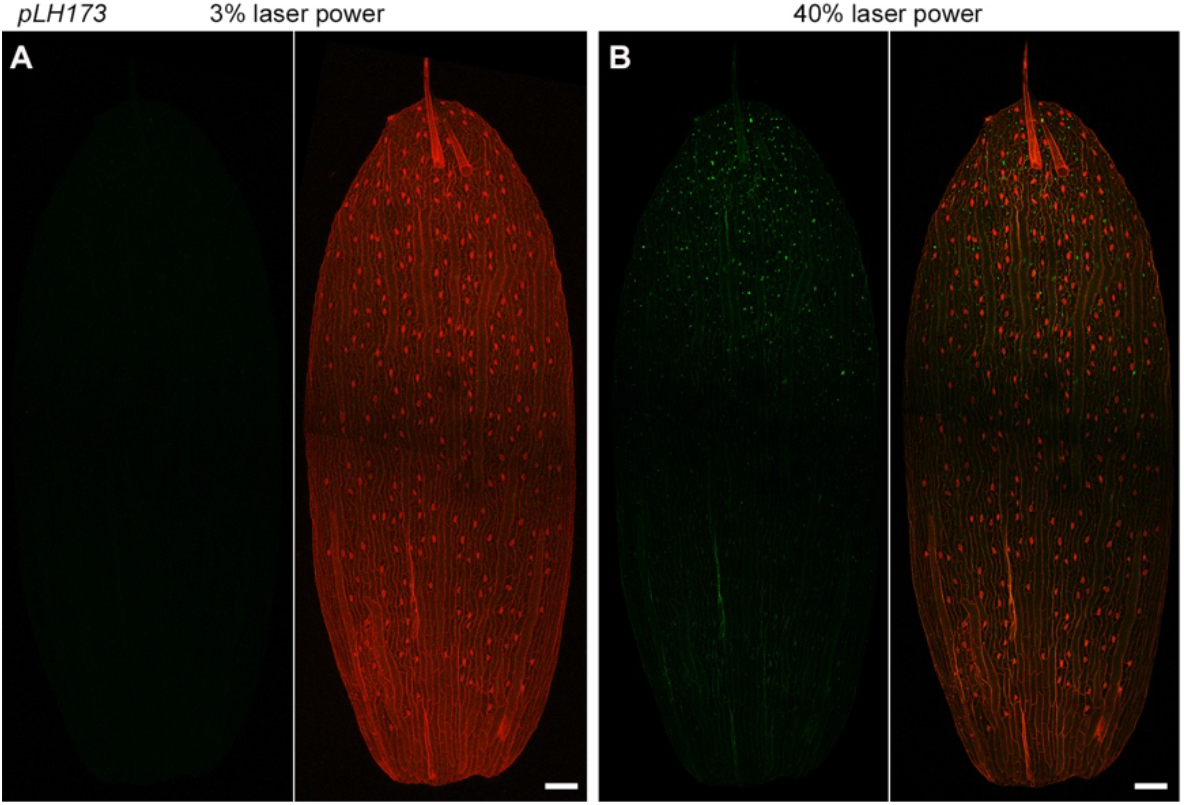
*pLH173* sepals expressed weak 3×Venus in some nuclei, related to Figure 3. Confocal images of a *pLH173* sepal imaged with 3% **(A)** and 40% **(B)** laser power. With 3% laser power, hardly any 3×Venus could be observed on *pLH173* sepal epidermis. With laser power increased to 40%, some nuclei at the sepal tip expressed weak 3×Venus. Images on the left, 3×Venus (green) marking the nuclei of cells expressing the enhancer reporter; Images on the right, 3×Venus (green) signals merged with PI (red) stained cell wall signals. A was duplicated from Fig.3D for comparison.

**Figure S6.**
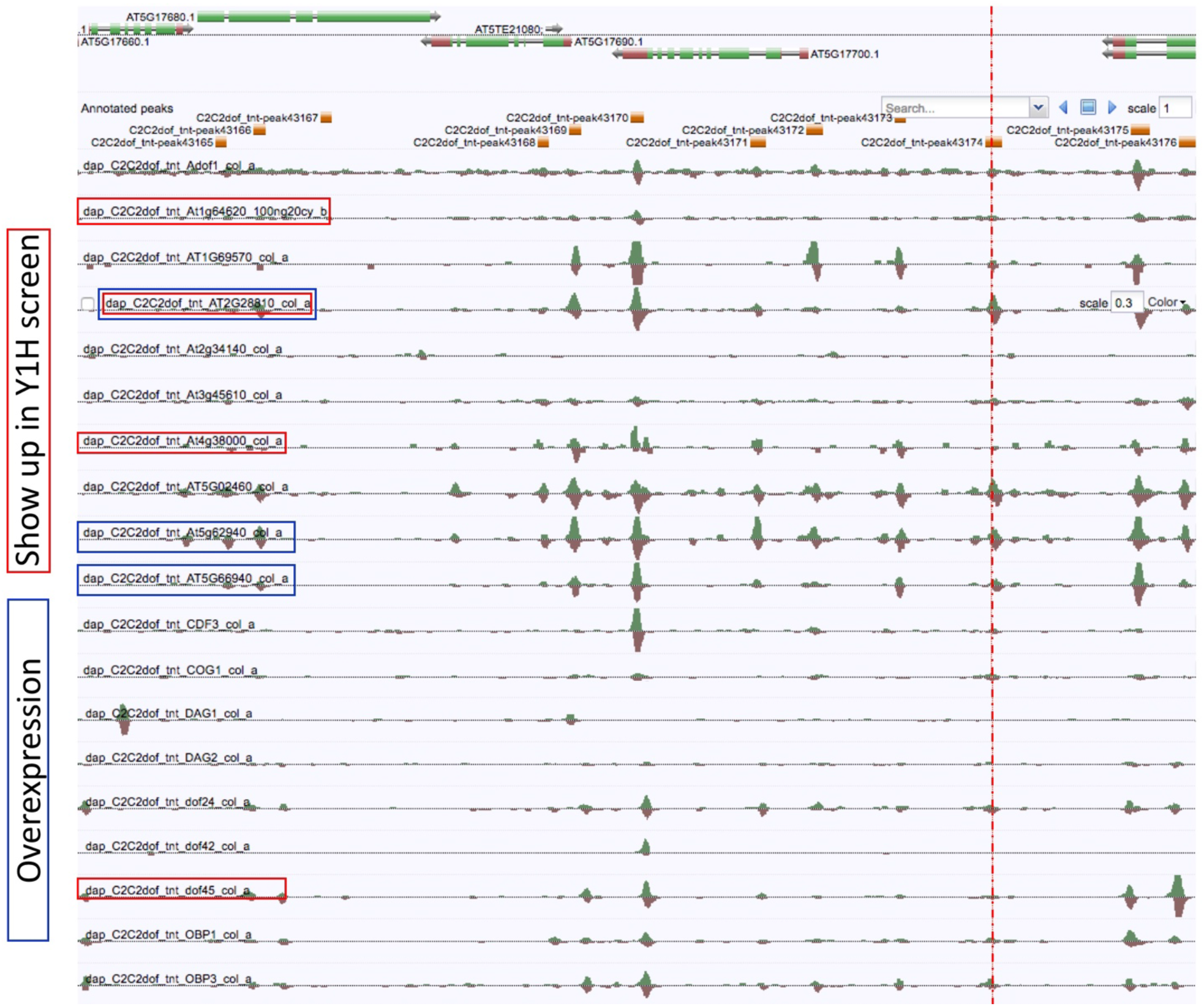
Binding sites of Dof transcription factors in the giant cell enhancer region and nearby, related to Figure 6. Results by DAP-seq. Dof transcription factors framed in red were identified in the yeast one-hybrid (Y1H) assay in this study; framed in blue were overexpressed in *pAR111* Arabidopsis. The red dash line marks the Dof binding site region.

**Figure S7.**
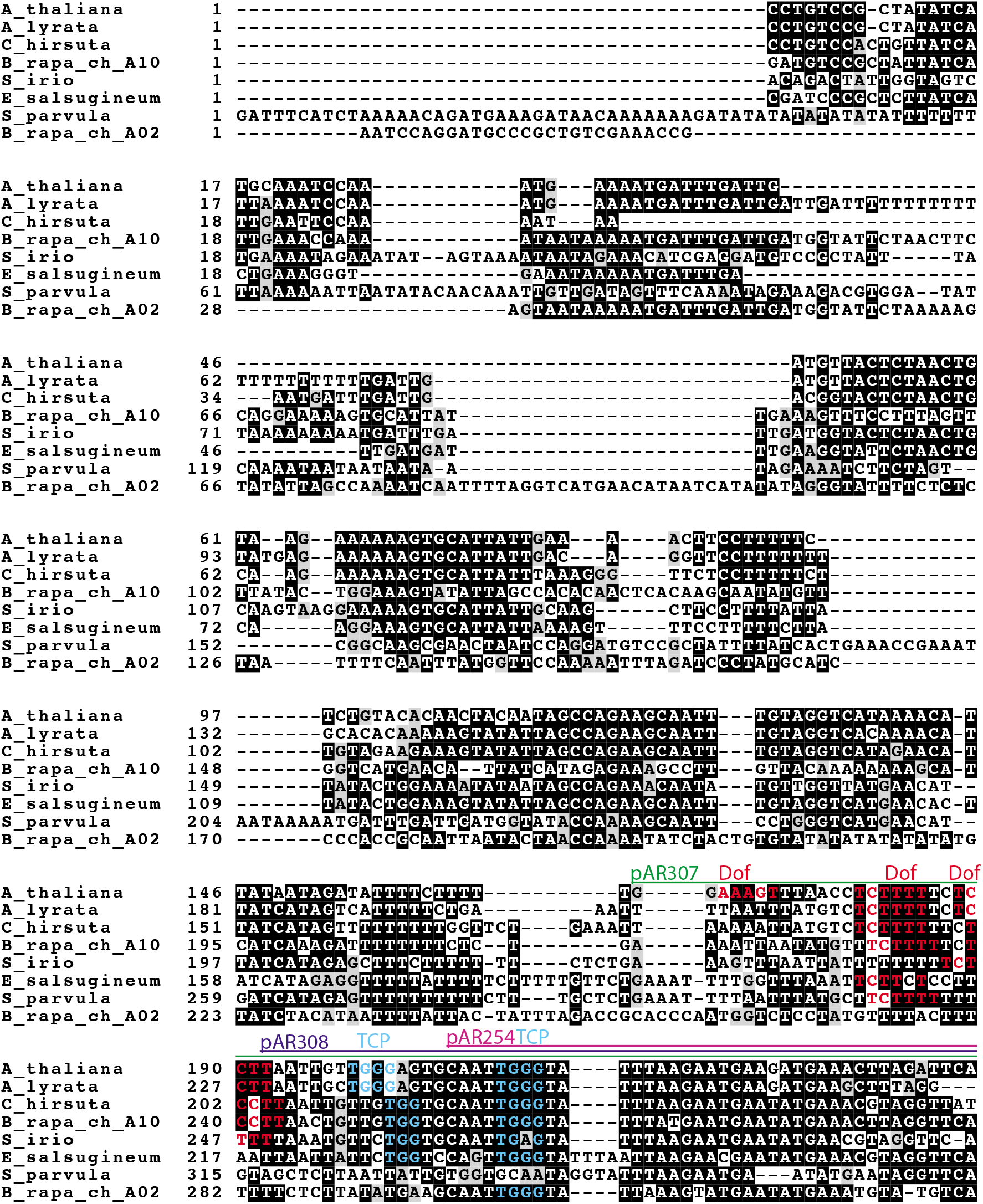
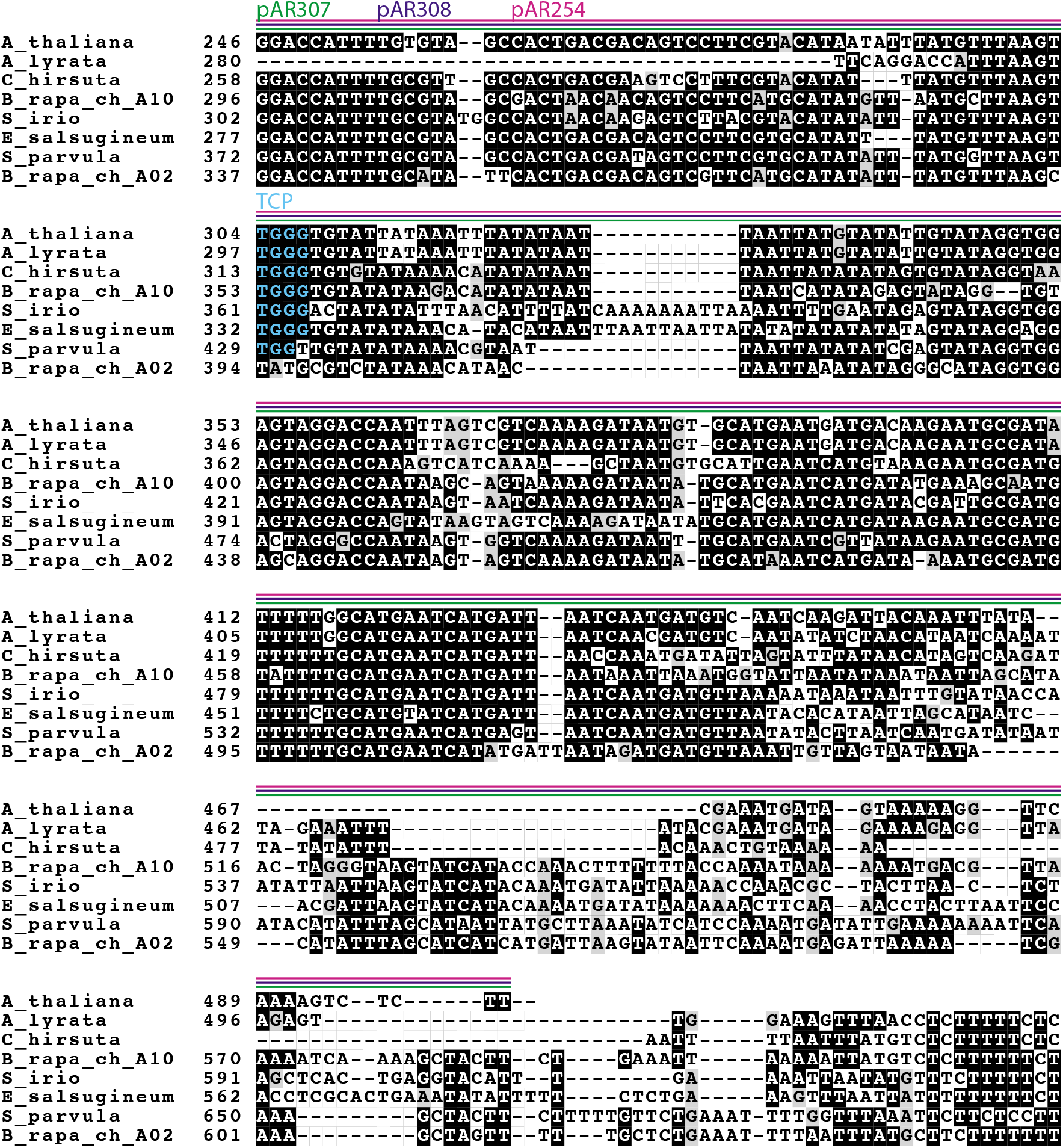
Conservation of Regions 1 and 2 of the giant cell enhancer across Brassicaceae species, related to Figure 4. Clustal omega sequence alignment of the giant cell enhancer Region 1-2 from *Arabidopsis thaliana* with *Arabidopsis lyrata, Cardamine hirsuta, Sisimbrium irio, Eutrema salsugineum, Schrenkiella parvula* (previously *Thellungiella parvula*) and *Brassica rapa*. Although *Brassica rapa* is a diploid, it underwent genome duplication during evolution, so there are two regions on chromosomes A10 and A02 which both align to the giant cell enhancer. The sequences included in *pAR307* (Region 2 + Dof, green line), *pAR308* (Region 2 plus, purple line), and *pAR254* (Region 2 only, magenta) are annotated with lines above the sequence. The putative Dof binding sites predicted in *Arabidopsis thaliana* and their conserved counterparts are highlighted with red letters. The putative TCP binding sites and their conserved counterparts are highlighted with aqua letters.

**Table S2:** Giant cell enhancer dissection constructs.

| **Plant Construct** | **Region of enhancer** | **Forward primer** | **Reverse primer** | **Entry clone** |
| --- | --- | --- | --- | --- |
| pAR111 | Full length (Region 1-2-3-4)<br>1-1024 bp | oAR214 | oAR215 | NA (Roeder et al., 2012) |
| pAR262 | Region 1-2-3<br>1-760 bp | oAR500 | oAR215 | pSL8 |
| pAR257 | Region 1-2<br>1-449 bp | oAR497 | oAR215 | pSL4 |
| pAR260 | Region 2-3-4<br>207-1024 bp | oAR214 | oAR498 | pSL7 |
| pAR261 | Region 2-3<br>207-760 bp | oAR500 | oAR498 | pSL8 |
| pAR254 | Region 2<br>207-499 bp | oAR497 | oAR498 | pSL1 |
| pAR258 | Region 3-4<br>450-1024 bp | oAR214 | oAR501 | pSL5 |
| pAR256 | Region 4<br>761-1024 bp | oAR214 | oAR499 | pSL3 |
| pAR280 | Region 1<br>1-208 bp | oAR532 | oAR215 | pAR278 |
| pLH173 | Region 2+ with 3 putative TCP sites mutated (oAR630-oAR631 used to mutate third TCP site)<br>190 – 449 bp | oAR497 | oAR612 | pDR9 |
| pAR307 | Region2 with Dof region<br>167-449 bp | oAR497 | oAR556 | pAR299 |
| pLH166 | Region 2 with 3 putative Dof sites mutated<br>167-449 bp | oAR497 | oLH412 | pLH164 |
| pAR308 | Region 2+ (without Dof region) 190 – 449 bp | oAR497 | oAR609 | pAR301 |

Associated with Methods

**Table S3:**
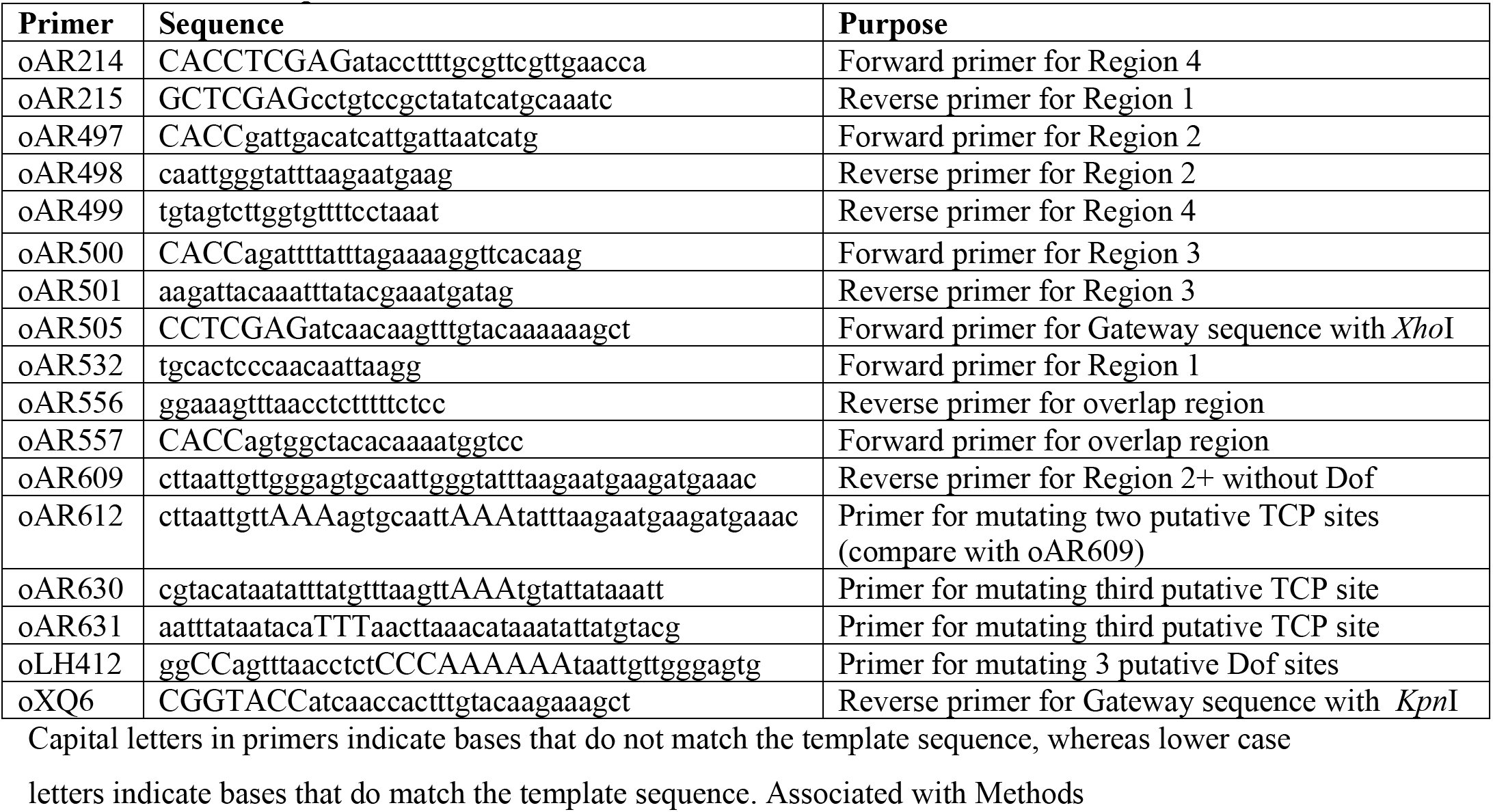
Primer sequences.

**Table S4:** Yeast one hybrid constructs.

| Yeast Construct | Region of enhancer | Forward primer | Reverse primer | Entry clone |
| --- | --- | --- | --- | --- |
| pAR276 | Region 1<br>1-208 bp | oAR215 | oAR532 | pAR278 |
| pAR275 | Region 2<br>207-499 bp | oAR498 | oAR497 | pSL1 |
| pAR277 | Overlap of junction<br>207-499 bp | oAR557 | oAR556 | pAR279 |

Associated with Methods

